# Neurotrophic factor Neuritin modulates T cell electrical and metabolic state for the balance of tolerance and immunity

**DOI:** 10.1101/2024.01.31.578284

**Authors:** Hong Yu, Hiroshi Nishio, Joseph Barbi, Marisa Mitchell-Flack, Paolo D. A. Vignali, Ying Zheng, Andriana Lebid, Kwang-Yu Chang, Juan Fu, Makenzie Higgins, Ching-Tai Huang, Xuehong Zhang, Zhiguang Li, Lee Blosser, Ada Tam, Charles G. Drake, Drew M. Pardoll

**Author notes:** These authors contributed equally to this work (Hong Yu, Hiroshi Nishio, Joseph Barbi).

## Abstract

The adaptive T cell response is accompanied by continuous rewiring of the T cell’s electric and metabolic state. Ion channels and nutrient transporters integrate bioelectric and biochemical signals from the environment, setting cellular electric and metabolic states. Divergent electric and metabolic states contribute to T cell immunity or tolerance. Here, we report that neuritin (Nrn1) contributes to tolerance development by modulating regulatory and effector T cell function. Nrn1 expression in regulatory T cells promotes its expansion and suppression function, while expression in the T effector cell dampens its inflammatory response. Nrn1 deficiency causes dysregulation of ion channel and nutrient transporter expression in Treg and effector T cells, resulting in divergent metabolic outcomes and impacting autoimmune disease progression and recovery. These findings identify a novel immune function of the neurotrophic factor Nrn1 in regulating the T cell metabolic state in a cell context-dependent manner and modulating the outcome of an immune response.

## Introduction

Peripheral T cell tolerance is important in restricting autoimmunity and minimizing collateral damage during active immune reactions and is achieved via diverse mechanisms, including T cell anergy, regulatory T (Treg) cell mediated suppression, and effector T (Te) cell exhaustion or deletion (ElTanbouly and Noelle, 2021). Upon activation, Treg and conventional T cells integrate environmental cues and adapt their metabolism to the energetic and biosynthetic demands, leading to tolerance or immunity. Tolerized versus responsive T cells are characterized by differential metabolic states. For example, T cell anergy is associated with reduced glycolysis, whereas activated T effector cells exhibit increased glycolysis (Buck et al., 2017; Geltink et al., 2018; Peng and Li, 2023; Zheng et al., 2009). Cellular metabolic states depend on electrolyte and nutrient uptake from the microenvironment (Chapman and Chi, 2022; Olenchock et al., 2017). Ion channels and nutrient transporters, which can integrate environmental nutrient changes, affect the cellular metabolic choices and impact the T cell functional outcome (Babst, 2020; Bohmwald et al., 2021; Ramirez et al., 2018). Each cell’s functional state would correspond with a set of ion channels and nutrient transporters supporting their underlying metabolic requirements. The mechanisms coordinating the ion channel and nutrient transporter expression changes to support the adaptive T cell functional state in the immune response microenvironment remain unclear.

Nrn1, also known as candidate plasticity gene 15 (CPG15), was initially discovered as a neurotrophic factor linked to the neuronal cell membrane through a glycosylphosphatidylinositol (GPI) anchor (Nedivi et al., 1998; Zhou and Zhou, 2014). It is highly conserved across species, with 98% overall homology between the murine and human protein. Nrn1 plays multiple roles in neural development, synaptic plasticity, synaptic maturation, neuronal migration, and survival (Cantallops et al., 2000; Javaherian and Cline, 2005; Nedivi et al., 1998; Putz et al., 2005; Zito et al., 2014). In the immune system, Nrn1 expression has been found in Foxp3^+^ Treg and follicular regulatory T cells (Tfr) (Gonzalez-Figueroa et al., 2021; Vahl et al., 2014), T cells from transplant tolerant recipients (Lim et al., 2013), anergized CD8 cells or CD8 cells from tumor-infiltrating lymphocytes in mouse tumor models (Schietinger et al., 2012; Schietinger et al., 2016; Singer et al., 2016), and in human Treg infiltrating breast cancer tumor tissue (Plitas et al., 2016). Soluble Nrn1 can be released from Tfr cells and act directly on B cells to suppress autoantibody development against tissue-specific antigens (Gonzalez-Figueroa et al., 2021). Despite the observation of Nrn1 expression in Treg cells and T cells from tolerant environments (Gonzalez-Figueroa et al., 2021; Lim et al., 2013; Plitas et al., 2016; Schietinger et al., 2012; Schietinger et al., 2016; Singer et al., 2016), the roles of Nrn1 in T cell tolerance development and Treg cell function have not been explored, and functional mechanism of Nrn1 remains elusive. This study demonstrates that the neurotrophic factor Nrn1 can moderate T cell tolerance and immunity through both Treg and Te cells, impacting Treg cell expansion and suppression while controlling inflammatory response in Te cells.

## Results

### Nrn1 expression and function in T cell anergy

To explore the molecular mechanisms underlying peripheral tolerance development, we utilized a system we previously developed to identify tolerance-associated genes (Huang et al., 2004). We compared the gene expression patterns associated with either a T effector/memory response or tolerance induction triggered by the same antigen but under divergent *in vivo* conditions (Huang et al., 2004). Influenza hemagglutinin (HA) antigen-specific TCR transgenic CD4 T cells were adoptively transferred into WT recipients with subsequent HA-Vaccinia virus (VacHA) infection to generate T effector/memory cells while tolerogenic HA-specific CD4s were generated by transfer into hosts with transgenic expression of HA as self-antigen (C3-HA mice, Figure 1A.)(Huang et al., 2004). One of the most differentially expressed genes upregulated in the anergy-inducing condition was Nrn1. Nrn1 expression was significantly higher among cells recovered from C3-HA hosts vs. cells from VacHA infected mice at all time points tested by qRT-PCR (Figure 1A). To further confirm the association of Nrn1 expression with T cell anergy, we assessed Nrn1 expression in naturally occurring anergic polyclonal CD4^+^ T cells (Ta), which can be identified by surface co-expression of Folate Receptor 4 (FR4) and the ecto-5’-nucleotidase CD73 (Ta, CD4^+^CD44^+^FR4^hi^CD73^hi^ cells)(Kalekar et al., 2016). Nrn1 expression was significantly higher in Ta than in naïve CD4 (Tn, CD4^+^CD62L^+^CD44^-^FR4^-^CD73^-^) and antigen-experienced cells (Te, CD4^+^CD44^+^FR4^-^CD73^-^) under steady-state conditions measured by both qRT-PCR and western blot (Figure 1B). Given that Treg cells, like anergic cells, have roles in maintaining immune tolerance, we queried whether Nrn1 is also expressed in Treg cells. Nrn1 expression can be detected in nTreg and induced Treg (iTreg) cells generated *in vitro* (Figure 1C). To evaluate Nrn1 expression under pathological tolerant conditions (Cuenca et al., 2003), we evaluated Nrn1 expression in T cells within the tumor microenvironment. Nrn1 expression in murine Treg cells and non-Treg CD4^+^ cells from tumor infiltrates were compared to the Treg cells and non-Treg CD4^+^ T cells isolated from peripheral blood. Nrn1 mRNA level was significantly increased among tumor-associated Treg cells and non-Treg CD4 cells compared to cells from peripheral blood (Figure 1-figure supplement 1A). Consistent with our findings in the mouse tumor setting, the Treg and non-Treg T cells from human breast cancer infiltrates reveal significantly higher Nrn1 expression compared to the peripheral blood Treg and non-Treg cells (Figure 1-figure supplement 1B) (Plitas et al., 2016).

**Figure 1.**
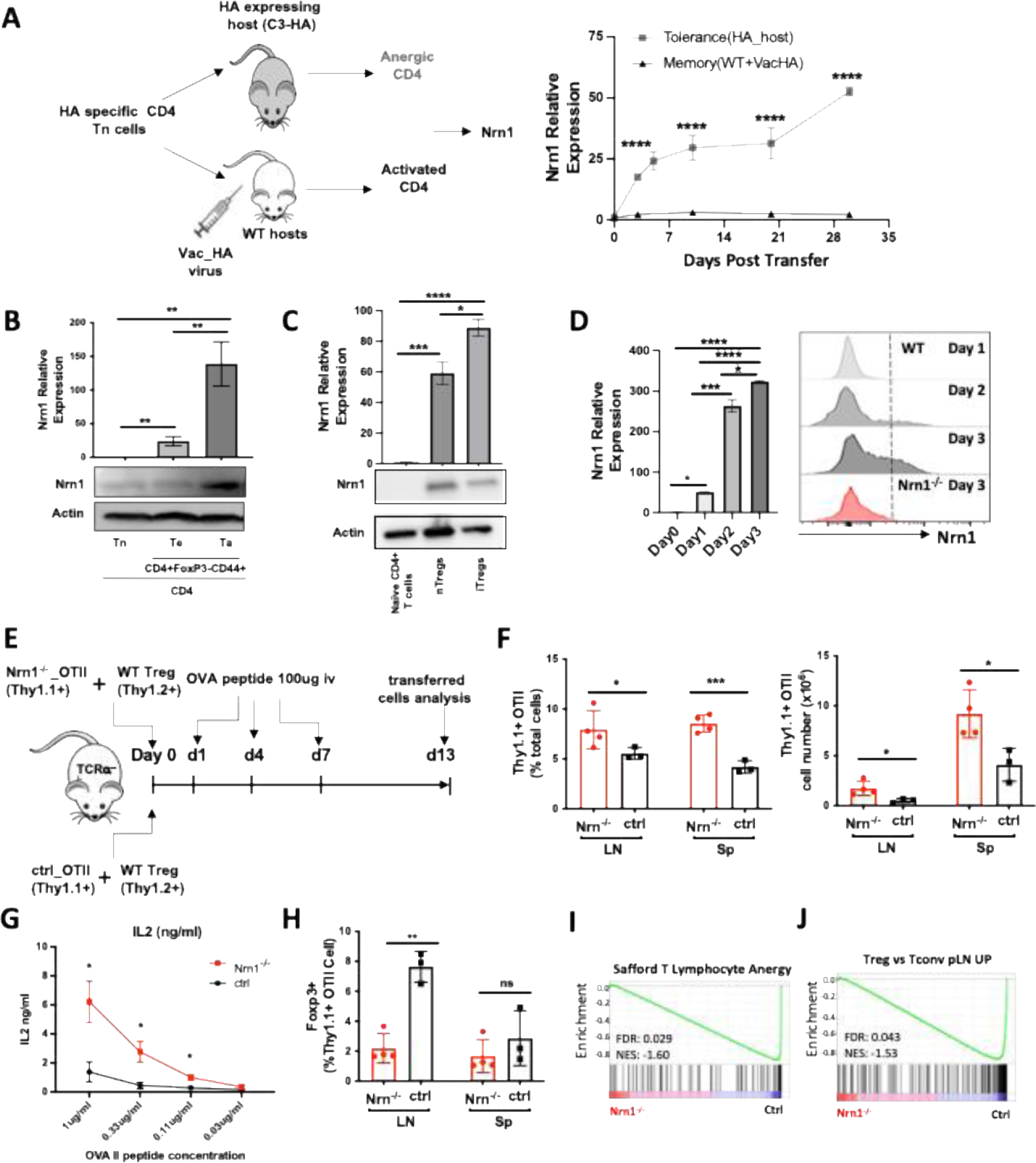
Nrn1 expression and function in anergic T cells. (**A**) Experimental scheme identifying Nrn1 in anergic T cells and qRT-PCR confirmation of Nrn1 expression in HA-specific CD4 cells recovered from HA-expressing host vs WT host activated with Vac_HA virus. (**B**) qRT-PCR and western blot detecting Nrn1 expression in naïve CD4^+^CD62L^hi^CD44^lo^ Tn cell, CD4 effector CD4^+^Foxp3^-^CD44^hi^CD73^-^FR^-^ Te cells and CD4 anergic CD4^+^Foxp3^-^CD44^hi^CD73^+^FR^+^ Ta cells. (**C**) Nrn1 expression was measured by qRT-PCR and western blot among naïve CD4^+^ T cells, CD4^+^Foxp3^+^ nTreg, and *in vitro* generated iTregs. (**D**) Nrn1 expression was detected by qRT-PCR and flow cytometry among WT naïve CD4^+^ cells and activated CD4^+^ cells on days 1, 2, and 3 after activation. Nrn1^-/-^ CD4 cells were also stained for Nrn1 three days after activation. qPCR Data are presented as average ± SEM. *p<0.05, **p<0.01, ***p<0.001, ****p<0.0001. Triplicates were used. Ordinary one-way ANOVA was performed for multi-comparison. (**E-J**). Anergy induction *in vivo*. (**E**) Experimental outline evaluating anergy development *in vivo*: 2x10^6^ Thy1.1^+^ Nrn1^-/-^ or ctrl CD4 OTII T cells were co-transferred with 5x10^5^ Thy1.2^+^Thy1.1^-^ WT Treg cells into TCRα^-/-^mice. Cells were recovered on day 13 post-transfer. (**F**) Proportions and numbers of OTII cells recovered from recipient spleen; (**G**) IL2 secretion from OTII cells upon *ex vivo* stimulation with OVA peptide. (**H**) Foxp3^+^ cell proportion among Thy1.1^+^ Nrn1^-/-^ or ctrl CD4 cells. (**I & J**) Nrn1^-/-^ vs ctrl OTII cells recovered from the peptide-induced anergy model were subjected to bulk RNASeq analysis. GSEA comparing the expression of signature genes for anergy (**I**) and Treg (**J**) among ctrl and Nrn1^-/-^ OTII cells. Data are presented as mean ±SEM and representative of 3 independent experiments (N≥4 mice per group). *p<0.05, **p<0.01, ***p<0.001. Unpaired Student’s t-tests were performed.

CD4^+^ T cells may pass through an effector stage after activation before reaching an anergic state (Adler et al., 1998; Chen et al., 2004; Huang et al., 2003; Opejin et al., 2020). To evaluate the potential role of Nrn1 expression in T cell tolerance development, we further examined Nrn1 expression kinetics after T cell activation. Nrn1 expression was significantly induced after CD4^+^ T cell activation (Figure 1D). Using an Nrn1-specific, monoclonal antibody, Nrn1 can be detected on activated CD4^+^ and CD8^+^ cells (Figure 1D, Figure 1-figure supplement 1C). The significant enhancement of Nrn1 expression after T cell activation suggests that Nrn1 may contribute to the process of T cell tolerance development and/or maintenance. Although Treg cells express Nrn1, we were not able to consistently detect substantial cell surface Nrn1 expression (Figure 1D, Figure 1-figure supplement 1D), likely due to Nrn1 being produced in a soluble form or cleaved from the cell membrane (Gonzalez-Figueroa et al., 2021).

To understand the functional implication of Nrn1 expression in immune tolerance, we analyzed Nrn1-deficient (Nrn1^-/-^) mice (Fujino et al., 2011). In the first evaluation of the Nrn1^-/-^ colony, Nrn1^-/-^ mice had consistently reduced body weight compared to heterozygous Nrn1^+/-^ and WT (Nrn1^+/+^) mice (Figure 1-figure supplement 2A). The lymphoid tissues of Nrn1^-/-^ mice were comparable to their Nrn1^+/-^ and WT counterparts except for a slight reduction in cell number that was observed in the spleens of Nrn1^-/-^ mice, likely due to their smaller size (Figure 1-figure supplement 2B). Analysis of thymocytes revealed no defect in T cell development (Figure 1-figure supplement 2C), and a flow cytometric survey of the major immune cell populations in the peripheral lymphoid tissue of these mice revealed similar proportions of CD4, CD8 T cells, B cells, monocytes and dendritic cells (DCs) (Figure 1-figure supplement 2D). Similarly, no differences were found between the proportions of anergic and Treg cells in Nrn1^-/-^, Nrn1^+/-^ and WT mice (Figure 1-figure supplement 2E, F), suggesting that Nrn1 deficiency does not significantly affect anergic and Treg cell balance under steady state. Additionally, histopathology assessment of lung, heart, liver, kidney, intestine and spleen harvested from 13 months old Nrn1^-/-^ and Nrn1^+/-^ littermates did not reveal any evidence of autoimmunity (data not shown). The comparable level of anergic and Treg cell population among Nrn1^-/-^, Nrn1^+/-^ and WT mice and lack of autoimmunity in Nrn1^-/-^ aged mice suggest that Nrn1 deficiency is not associated with baseline immune abnormalities or overt dysfunction. Due to the similarity between Nrn1^+/-^ and WT mice, we have used either Nrn1^+/-^ or WT mice as our control depending on mice availability and referred to both as “ctrl” in the subsequent discussion.

To evaluate the relevance of Nrn1 in CD4^+^ T cell tolerance development, we employed the classic peptide-induced T cell anergy model (Vanasek et al., 2006). Specifically, we crossed OVA antigen-specific TCR transgenic OTII mice onto the Nrn1^-/-^ background. Nrn1^-/-^_OTII^+^ or control_OTII^+^ (ctrl_OTII^+^) cells marked with Thy1.1^+^ congenic marker (Thy1.1^+^Thy1.2^-^), were co-transferred with polyclonal WT Tregs (marked as Thy1.1^-^thy1.2^+^), into TCRα knockout mice (TCRα^-/-^), followed by injection of soluble OVA peptide to induce clonal anergy (Figure 1E) (Chappert and Schwartz, 2010; Martinez et al., 2012; Mercadante and Lorenz, 2016; Shin et al., 2014). On day 13 after cell transfer, the proportion and number of OTII cells increased in the Nrn1^-/-^_OTII compared to the ctrl_OTII hosts (Figure 1F). Moreover, Nrn1^-/-^_OTII cells produced increased IL2 than ctrl_OTII upon restimulation (Figure 1G). Anergic CD4 Tconv cells can transdifferentiate into Foxp3^+^ pTreg cells *in vivo* (DL, 2017; Kalekar et al., 2016; Kuczma et al., 2021). Consistent with reduced anergy induction, the proportion of Foxp3^+^ pTreg among Nrn1^-/-^_OTII was significantly reduced (Figure 1H). In parallel with the phenotypic analysis, we compared gene expression between Nrn1^-/-^_OTII and ctrl_OTII cells by RNA Sequencing (RNASeq). Gene set enrichment analysis (GSEA) revealed that the gene set on T cell anergy was enriched in ctrl relative to Nrn1^-/-^_OTII cells (Figure 1I)(Safford et al., 2005). Also, consistent with the decreased transdifferentiation to Foxp3^+^ cells, the Treg signature gene set was prominently reduced in Nrn1^-/-^_OTII cells relative to the ctrl (Figure 1J). Anergic T cells are characterized by inhibition of proliferation and compromised effector cytokines such as IL2 production (Choi and Schwartz, 2007). The increased cell expansion and cytokine production in Nrn1^-/-^_OTII cells and the reduced expression of anergic and Treg signature genes all support the notion that Nrn1 is involved in T cell anergy development.

Anergic T cells are developed after encountering antigen, passing through a brief effector stage, and reaching an anergic state (Chappert and Schwartz, 2010; Huang et al., 2003; Silva Morales and Mueller, 2018; Zha et al., 2006). Enhanced T cell activation, defective Treg cell conversion or expansion, and heightened T effector cell response may all contribute to defects in T cell anergy induction and/or maintenance (Chappert and Schwartz, 2010; Huang et al., 2003; Kalekar et al., 2016; Silva Morales and Mueller, 2018; Zha et al., 2006). We first examined early T cell activation to understand the underlying cause of defective anergy development in Nrn1^-/-^ cells. Nrn1^-/-^ CD4^+^ cells showed reduced T cell activation, as evidenced by reduced CellTrace violet dye (CTV) dilution, activation marker expression, and Ca^++^ entry after TCR stimulation (Figure 1-figure supplement 3A, B, C). The reduced early T cell activation observed in Nrn1^-/-^ CD4 cells suggests that the compromised anergy development in Nrn1^-/-^_OTII cells was not caused by enhanced early T cell activation. The defective pTreg generation and/or enhanced effector T cell response may contribute to compromised anergy development.

### Compromised Treg expansion and suppression in the absence of Nrn1

The significant reduction of Foxp3^+^ pTreg among Nrn1^-/-^_OTII cells could be caused by the diminished conversion of Foxp3^-^ Tconv cells to pTreg and/or diminished Treg cell expansion and persistence. To understand the cause of pTreg reduction in Nrn1^-/-^_OTII cells (Figure 1H), we turned to the induced Treg (iTreg) differentiation system to evaluate the capability of Foxp3^+^ Treg development and expansion in Nrn1^-/-^ cells. Similar proportions of Foxp3^+^ cells were observed in Nrn1^-/-^ and ctrl cells under the iTreg culture condition (Figure 2A), suggesting that Nrn1 deficiency does not significantly impact Foxp3^+^ cell differentiation. To examine the capacity of iTreg expansion, Nrn1^-/-^ and ctrl iTreg cells were restimulated with anti-CD3, and we found reduced live cells over time in Nrn1^-/-^ iTreg compared to the ctrl (Figure 2B). The reduced live cell number in Nrn1^-/-^ was accompanied by reduced Ki67 expression (Figure 2C). Although Nrn1^-/-^ iTregs retained a higher proportion of Foxp3^+^ cells three days after restimulation, however, when taking into account the total number of live cells, the actual number of live Foxp3^+^ cells was reduced in Nrn1^-/-^ (Figure 2D). Treg cells are not stable and are prone to losing Foxp3 expression after extended proliferation (Feng et al., 2014; Floess et al., 2007; Li et al., 2014; Zheng et al., 2010). The increased proportion of Foxp3^+^ cells was consistent with reduced proliferation observed in Nrn1^-/-^ cells. Thus, Nrn1 deficiency can lead to reduced iTreg cell proliferation and persistence *in vitro*.

**Figure 2.**
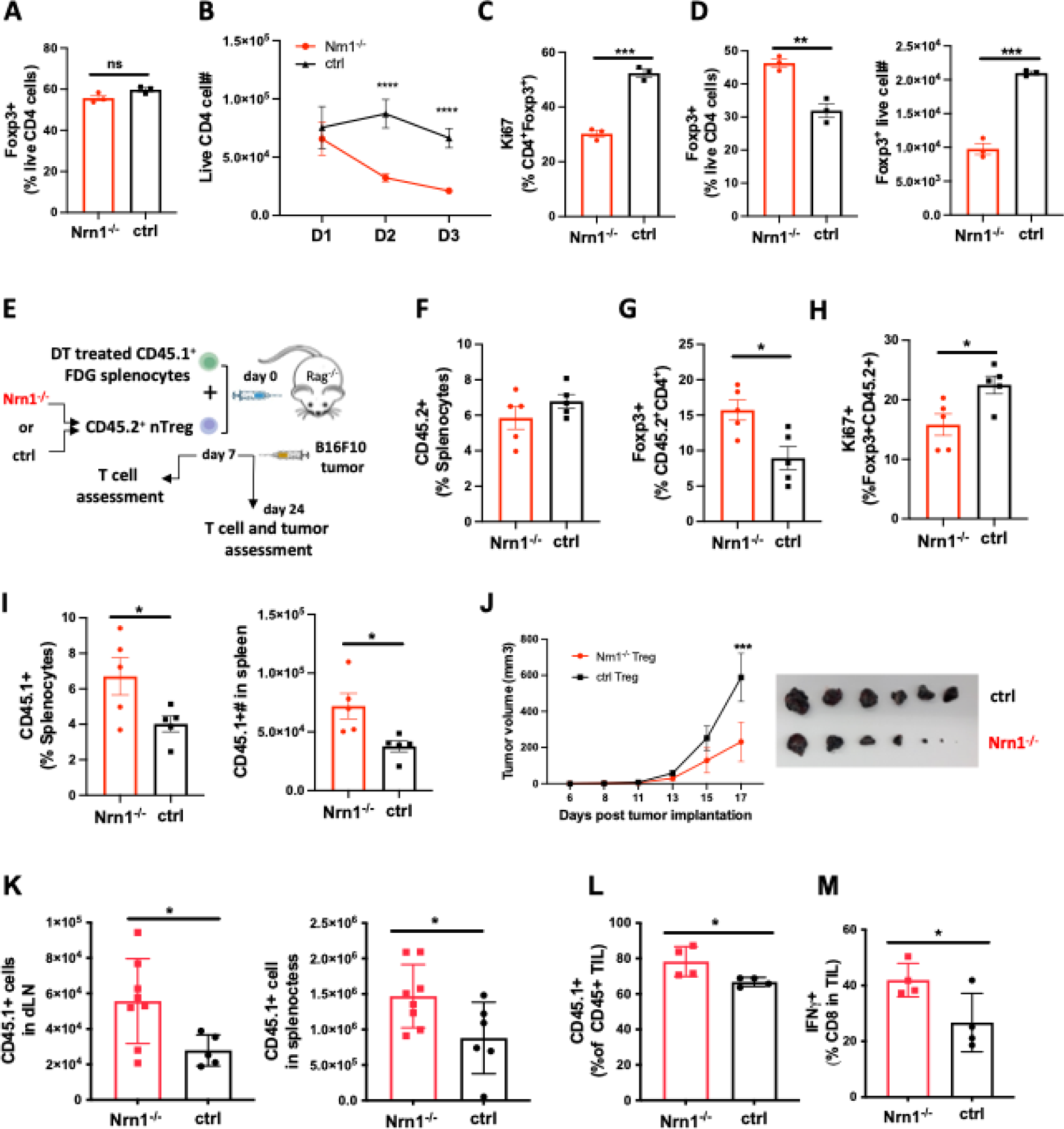
Reduced proliferation and suppression function in Nrn1^-/-^ Treg cells. (**A**). Proportion of Foxp3^+^ cells three days after *in vitro* iTreg differentiation. (**B-D**) iTreg cell expansion after restimulation. (**B**) The number of live cells from day 1 to day 3 after iTreg cell restimulation with anti-CD3. (**C**) Ki67 expression among CD4^+^Foxp3^+^ cells day 3 after restimulation. (**D**) Foxp3^+^ cell proportion and number among live CD4^+^ cells day 3 after restimulation. Triplicates in each experiment, data represent one of four independent experiments. (**E-M**). Nrn1^-/-^ or ctrl nTreg cells expansion and suppression *in vivo*. (**E**) The experimental scheme. CD45.2^+^ nTreg T cells from Nrn1^-/-^ or ctrl were transferred with CD45.1^+^ FDG splenocytes devoid of Tregs into Rag2^-/-^ host. Treg cell expansion and suppression toward FDG CD45.1^+^ responder cells were evaluated on day 7 post cell transfer. Alternatively, B16F10 tumor cells were inoculated on day 7 after cell transfer and monitored for tumor growth. (**F-J**) CD45.2^+^ cell proportion (**F**), Foxp3 retention (**G**), and Ki67 expression among Foxp3^+^ cells (**H**) at day 7 post cell transfer. (**I**) CD45.1^+^ cell proportion and number in the spleen of Nrn1^-/-^ or ctrl Treg hosts day 7 post cell transfer. (**J-L**). Treg cell suppression toward anti-tumor response. (**J**) Tumor growth curve and tumor size at harvest from Nrn1^-/-^ or ctrl nTreg hosts. (**K**) CD45.1^+^ cell count in tumor draining lymph node (LN) and spleen. (**L**) the proportion of CD45.1^+^ cells among CD45^+^ tumor lymphocyte infiltrates (TILs). (**M**) IFNγ% among CD8^+^ T cells in TILs. n≥5 mice per group. (**F-I**) represents three independent experiments, **(J-M**) represents two independent experiments. Data are presented as mean ±SEM *p<0.05, **p<0.01, ***p<0.001, ****p<0.0001. Unpaired Student’s t-tests were performed.

The defects observed in iTreg cell expansion *in vitro* prompt further examination of Nrn1^-/-^ nTreg expansion and suppression function *in vivo.* To this end, we tested the suppression capacity of congenically marked (CD45.1^-^CD45.2^+^) Nrn1^-/-^ or ctrl nTreg toward CD45.1^+^CD45.2^-^ responder cells in Rag2^-/-^ mice (Figure 2E). The CD45.1^+^CD45.2^-^ responder cells devoid of Treg cells were splenocytes derived from Foxp3DTRGFP (FDG) mice pretreated with diphtheria toxin (DT) (Kim et al., 2007; Workman et al., 2011). DT treatment caused the deletion of Treg cells in FDG mice (Kim et al., 2007). Although the CD45.1^-^CD45.2^+^ Nrn1^-/-^ and ctrl cell proportions were not significantly different among hosts splenocytes at day 7 post transfer (Figure 2F), Nrn1^-/-^ cells retained a higher Foxp3^+^ cell proportion and reduced Ki67 expression comparing to the ctrl (Figure 2G, H). These findings were similar to our observation of iTreg cells *in vitro* (Figure 2C, D). Nrn1^-/-^ Tregs also showed reduced suppression toward CD45.1^+^ responder cells, evidenced by increased CD45.1^+^ proportion and cell number in host splenocytes (Figure 2I).

To evaluate the functional implication of Nrn1^-/-^ Treg suppression in disease settings, we challenged the Rag2^-/-^ hosts with the poorly immunogenic B16F10 tumor (Figure 2E). Tumors grew much slower in Nrn1^-/-^ Treg recipients than those reconstituted with ctrl Tregs (Figure 2J). Moreover, the number of CD45.1^+^ cells in tumor-draining lymph nodes and spleens increased significantly in Nrn1^-/-^ Treg hosts compared to the ctrl group (Figure 2K). Consistently, the CD45.1^+^ responder cell proportion among tumor lymphocyte infiltrates (TILs) was also increased (Figure 2L), accompanied by an increased proportion of IFNγ^+^ cells among CD8 TILs from Nrn1^-/-^ Treg hosts (Figure 2M). The increased expansion of CD45.1^+^ responder cells and reduced tumor growth further confirmed the reduced suppressive capacity of Nrn1^-/-^ Treg cells.

### Nrn1 impacts Treg cell electrical and metabolic state

To understand the molecular mechanisms associated with Nrn1^-/-^ Treg cells, we compared gene expression between Nrn1^-/-^ and ctrl iTregs under resting (IL2 only) and activation (aCD3 and IL2) conditions by RNASeq. GSEA on gene ontology database and clustering of enriched gene sets by Cytoscape identified three clusters enriched in resting Nrn1^-/-^ iTreg (Figure 3A, Figure 3-figure supplement Table 1)(Shannon et al., 2003; Subramanian et al., 2005). The “neurotransmitter involved in membrane potential (MP)” and “sodium transport” clusters involved gene sets on the ion transport and cell MP regulation (Figure 3A, Figure 3-figure supplement Table 1). MP is the difference in electric charge between the interior and the exterior of the cell membrane (Abdul Kadir et al., 2018; Blackiston et al., 2009; Ma et al., 2017). Ion channels and transporters for Na^+^ and other ions such as K^+^, Cl^-^ *et al*. maintain the ion balance and contribute to cell MP (Blackiston et al., 2009). MP change can impact cell plasma membrane lipid dynamics and affect receptor kinase activity (Zhou et al., 2015). The enrichment of “receptor protein kinase” gene set clusters may reflect changes caused by MP (Figure 3A, Figure 3-figure supplement Table 1). Gene set cluster analysis on activated iTreg cells also revealed the enrichment of the “ion channel and receptor” cluster in Nrn1^-/-^ cells (Figure 3B, Figure 3-figure supplement table 2), supporting the potential role of Nrn1 in modulating ion balances and MP.

**Figure 3.**
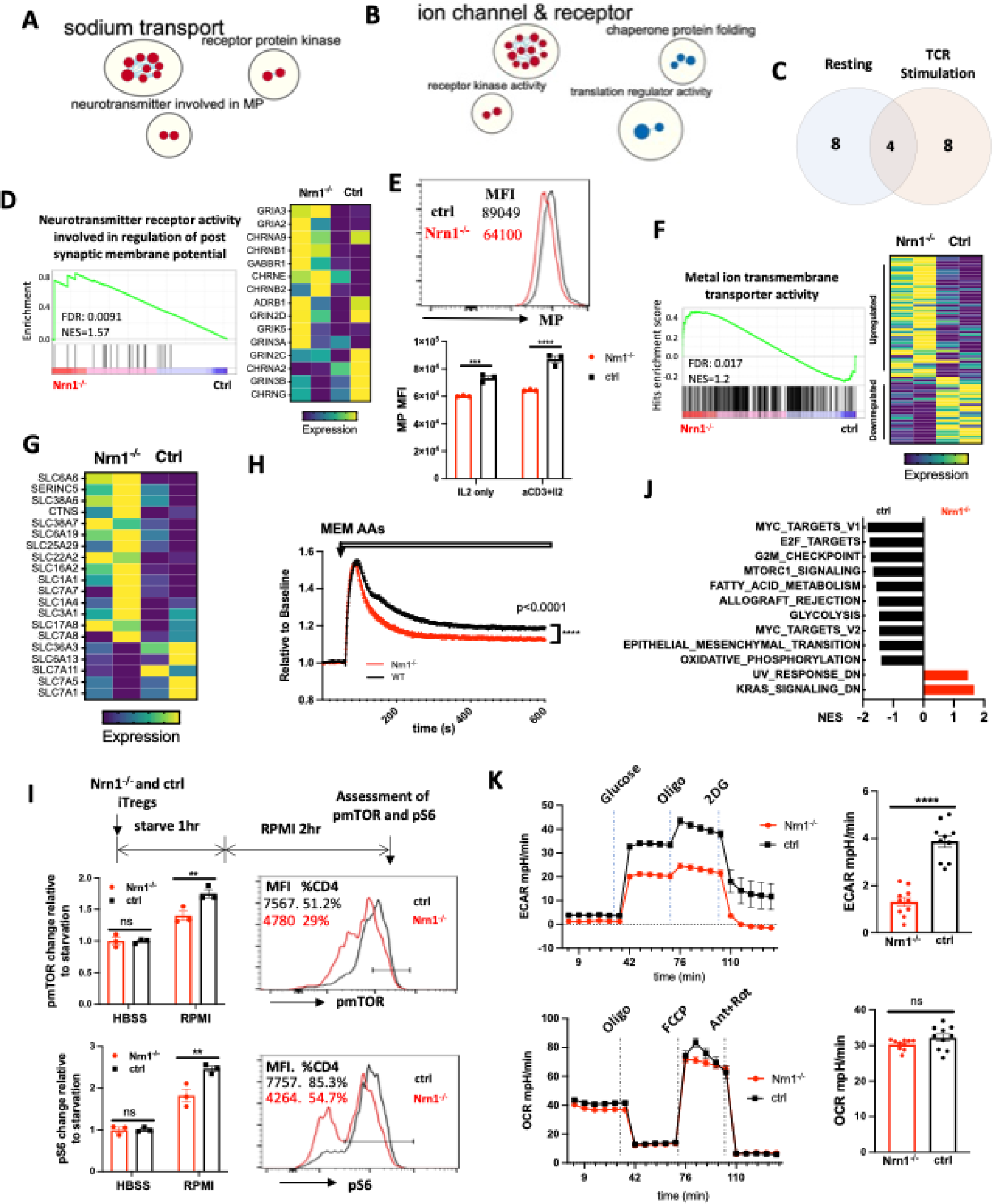
Nrn1 expression impacts Treg cell electrical and metabolic state. (**A-C**). Gene sets clusters enriched in Nrn1^-/-^ and ctrl iTreg cells. Gene sets cluster analysis via Cytoscape was performed on Gene ontology Molecular Function (GO_MF) gene sets. The results cutoff: p-value ≤0.05 and FDR q-value ≤0.1. (**A**) Gene sets cluster in Nrn1^-/-^ iTreg cells cultured under resting condition (IL2 only) (Figure 3-figure supplement Table 1). (**B**) Gene sets clusters in Nrn1^-/-^ and ctrl iTreg cells reactivated with anti-CD3 (Figure 3-figure supplement Table 2). (**C**) Comparison of enriched gene sets in Nrn1^-/-^ under resting vs. activating condition (Figure 3-figure supplement Table 3). (**D-F)** Changes relating to cell electric state. (**D**) Enrichment of “GOMF_Neurotransmitter receptor activity involved in the regulation of postsynaptic membrane potential” gene set and enriched gene expression heatmap. (**E**) Membrane potential was measured in Nrn1^-/-^ and ctrl iTreg cells cultured in IL2 or activated with anti-CD3 in the presence of IL2. Data represent three independent experiments. (**F**) Enrichment of “GOMF_Metal ion transmembrane transporter activity” gene set and enriched gene expression heatmap (Figure 3-figure supplement 1A). (**G-K**). Metabolic changes associated with Nrn1^-/-^ iTreg. (**G**) Heatmap of differentially expressed amino acid (AA) transport-related genes (from “MF_Amino acid transmembrane transporter activity” gene list) in Nrn1^-/-^ and ctrl iTreg cells. (**H**) AAs induced MP changes in Nrn1^-/-^ and ctrl iTreg cells. Data represent three independent experiments. (**I**) Measurement of pmTOR and pS6 in iTreg cells that were deprived of nutrients for 1h and refed with RPMI for two hours. (**J**) Hallmark gene sets significantly enriched in Nrn1^-/-^ and ctrl iTreg. NOM p-val<0.05, FDR q-val<0.25. (**K**) Seahorse analysis of extracellular acidification rate (ECAR) and oxygen consumption rate (OCR) in Nrn1^-/-^ and ctrl iTreg cells. n=6∼10 technical replicates per group. Data represent three independent experiments. **p<0.01, ***p<0.001, ****p<0.0001. Unpaired student t-test for two-group comparison. Unpaired t-test (H, K), two-way ANOVA (E, I). ns, not significant.

The “Neurotransmitter receptor activity involved in regulation of postsynaptic membrane potential” gene set was significantly enriched under resting and activation conditions (Figure 3C and D; Figure 3-figure supplement Table 3). The α-amino-3-hydroxy-5-methyl-4- isoxazolepropionic acid receptor (AMPAR) subunits Gria2 and Gria3 are the major components of this gene set and showed increased expression in Nrn1^-/-^ cells (Figure 3D). AMPAR is an ionotropic glutamate receptor that mediates fast excitatory synaptic transmission in neurons. Nrn1 has been reported as an accessory protein for AMPAR (Pandya et al., 2018; Schwenk et al., 2012; Subramanian et al., 2019), although the functional implication of Nrn1 as an AMPAR accessory protein remains unclear. The enrichment of MP related gene set prompted the examination of electric status, including MP level and ion channel expressions. We examined the relative MP level by FLIPR MP dye, a lipophilic dye able to cross the plasma membrane, which has been routinely used to measure cell MP changes (Dvorak et al., 2021; Joesch et al., 2008; Nik et al., 2017; Whiteaker et al., 2001). When the cells are depolarized, the dye enters the cells, causing an increase in fluorescence signal. Conversely, cellular hyperpolarization results in dye exit and decreased fluorescence. Compared to ctrl iTreg cells, Nrn1^-/-^ exhibits significant hyperpolarization under both resting and activation conditions (Figure 3E). Consistent with the MP change, the “MF_metal ion transmembrane transporter activity” gene set, which contains 436 ion channel related genes, was significantly enriched and showed a different expression pattern in Nrn1^-/-^ iTregs (Figure 3F; Figure 3-figure supplement 1A and B). The changes in cellular MP and differential expression of ion channel and transporter genes in Nrn1^-/-^ implicate the role of Nrn1 in the balance of electric state in the iTreg cell.

MP changes have been associated with changes in amino acid (AA) transporter expression and nutrient acquisition, which in turn influences cellular metabolic and functional state (Yu et al., 2022). To understand whether MP changes in Nrn1^-/-^ are associated with changes in nutrient acquisition and thus the metabolic state, we surveyed AA transport-related gene expression using the “Amino acid transmembrane transporter activity” gene set and found differential AA transporter gene expression between Nrn1^-/-^ and ctrl iTregs (Figure 3G). Upon AA entry through transporters, the electric charge carried by these molecules may transiently affect cell membrane potential. Differential AA transporter expression patterns may have different impacts on cellular MP upon AA entry. Thus, we loaded Nrn1^-/-^ and ctrl iTreg with FLIPR MP dye in the HBSS medium and tested cellular MP change upon exposure to MEM AAs. The AA-induced cellular MP change was reduced in Nrn1^-/-^ compared to the ctrl, reflective of differential AA transporter expression patterns (Figure 3H). Electrolytes and AAs entry are critical regulators of mTORC1 activation and T cell metabolism (Liu and Sabatini, 2020; Saravia et al., 2020; Sinclair et al., 2013; Wang et al., 2020). We examined mTORC1 activation at the protein level by evaluating mTOR and S6 phosphorylation via flow cytometry. We found reduced phosphorylation of mTOR and S6 in activated Nrn1^-/-^ iTreg cells (Figure 3-figure supplement 1C). We further performed a nutrient-sensing assay to evaluate the role of ion and nutrient entry in mTORC1 activation. Nrn1^-/-^ and ctrl iTreg cells were starved for one hour in a nutrient-free buffer, followed by adding RPMI medium with complete ions and nutrients, and cultured for two more hours. While adding the medium with nutrients clearly increased the mTOR and S6 phosphorylation, the degree of change was significantly less in Nrn1^-/-^ than in the ctrl (Figure 3I). Consistently, GSEA on Hallmark gene sets reveal reduced gene set enrichment relating to the mTORC1 signaling, corroborating the reduced pmTOR and pS6 detection in Nrn1^-/-^ cells. Moreover, Nrn1^-/-^ cells also showed reduced expression of glycolysis, fatty acid metabolism, and oxidative phosphorylation related gene sets under both resting and activating conditions (Figure 3J, Figure 3-figure supplement 1D), indicating changes in metabolic status. Since previous work has identified mTORC1 to be an important regulator of aerobic glycolysis and given that our GSEA data suggested changes in glycolysis (Figure 3J) (Salmond, 2018), we performed the seahorse assay and confirmed reduced glycolysis among Nrn1^-/-^ cells (Figure 3K). Examination of mitochondrial bioenergetic function revealed a similar oxygen consumption rate (OCR) between Nrn1^-/-^ and ctrl cells (Figure 3K). Thus, Nrn1 expression can affect the iTreg electric state, influence ion channel and nutrient transporter expression, impact nutrient sensing, modulate metabolic state, and contribute to Treg expansion and suppression function.

We have observed significant changes in the electrical and metabolic state among Nrn1^-/-^ iTreg compared to the ctrl. Because Nrn1 can be expressed on the cell surface, one question arises whether the changes observed in Nrn1^-/-^ cells were caused by the functional deficiency of Nrn1 or arose secondary to potential changes in cell membrane structure originating at the Nrn1^-/-^ naïve T cell stage. To answer this question, we first examined potential changes in electrical and metabolic status among Nrn1^-/-^ naïve CD4 T cells. The Nrn1^-/-^ naïve CD4 T cells showed similar resting MP and AA-induced MP changes compared to the ctrl cells (Figure 3-supplement figure 2 A, B). We also observed comparable glycolysis and mitochondrial bioenergetic function between Nrn1^-/-^ naïve CD4 T cells and the ctrl (Figure 3-supplement figure 2 C). These results suggest the electrical and metabolic state in Nrn1^-/-^ T cells are comparable to the ctrl cells at the naïve cell stage. To further rule out the possibility that the observed changes in Nrn1^-/-^ iTreg are secondary to developmental structural changes, not Nrn1 functional deficiency, we differentiated WT T cells in the presence of antagonistic Nrn1 antibody and compared to the WT ctrl and Nrn1^-/-^ iTreg cells. WT iTreg cells differentiated in the presence of Nrn1 antibody exhibit reduced resting MP, similar to Nrn1^-/-^ cells (Figure 3-supplement figure 2 D). Moreover, upon restimulation, WT iTreg cells differentiated under Nrn1 antibody blockade showed a similar phenotype as Nrn1^-/-^ cells, with reduced live cell number, reduced Ki67 expression, and increased Foxp3^+^ cell proportion among live cells (Figure 3-supplement figure 2 E, F). These results suggest that Nrn1 functional deficiency likely contributes to the electrical and metabolic state change observed in Nrn1^-/-^ iTreg cells.

### Nrn1 impact effector T cell inflammatory response

CD4^+^ T cells can pass through an effector stage on their way to an anergic state (Huang et al., 2003). Since Nrn1 expression is significantly induced after T cell activation (Figure 1D), Nrn1 might influence CD4^+^ effector (Te) cell differentiation, affecting anergy development. Nrn1 may exert different electric changes due to distinct ion channel expression contexts in Te cells than in Tregs. We first evaluated Nrn1^-/-^ Te cell differentiation *in vitro.* Nrn1 deficient CD4 Te cells showed increased Ki67 expression, associated with increased cytokine TNFα, IL2, and IFNγ expression upon restimulation (Figure 4A). To evaluate Nrn1^-/-^ Te cell response *in vivo*, we crossed Nrn1^-/-^ with FDG mice and generated Nrn1^-/-^_FDG and ctrl_FDG mice, which enabled the elimination of endogenous Treg cells (Figure 4B). Deleting endogenous Foxp3^+^ Treg cells using DT will cause the activation of self-reactive T cells, leading to an autoimmune response (Kim et al., 2007; Nystrom et al., 2014). Upon administration of DT, we observed accelerated weight loss in Nrn1^-/-^_FDG mice, reflecting enhanced autoimmune inflammation (Figure 4C). Examination of T cell response revealed a significant increase in Ki67 expression and inflammatory cytokine TNFα, IL2, and IFNγ expression among Nrn1^-/-^ CD4 cells on day 6 post DT treatment (Figure 4D), consistent with the findings *in vitro*. The proportion of Foxp3^+^ cells was very low on day 6 post DT treatment and comparable between Nrn1^-/-^ and the ctrl (Figure 4E), suggesting that the differential Te cell response was not due to the impact from Treg cells. Thus, Nrn1 deficiency enhances Te cell response *in vitro* and *in vivo*.

**Figure 4.**
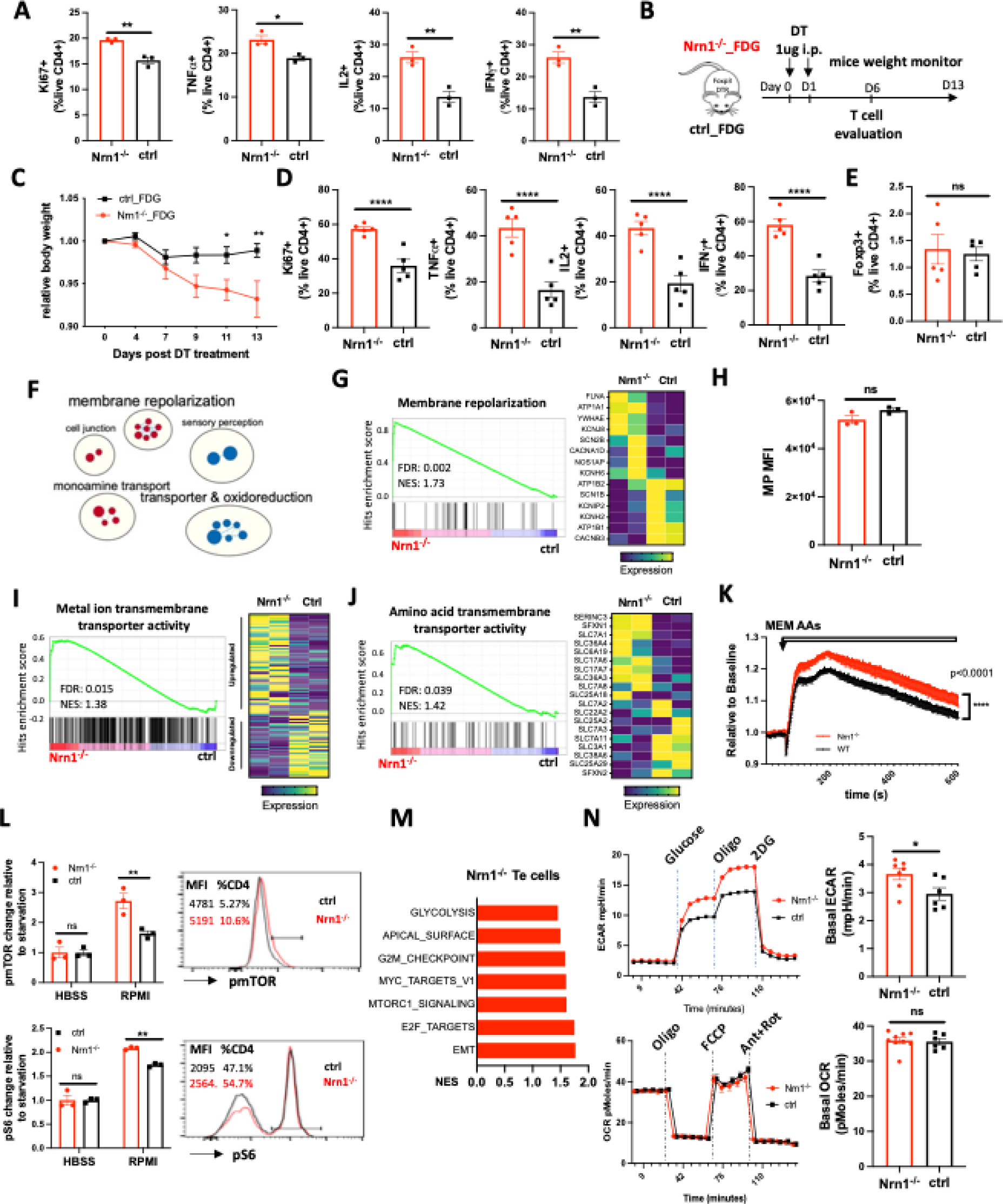
Nrn1 deficiency affects Te cell response. (**A**) Comparison of cell proliferation and cytokine expression in Nrn1^-/-^ and ctrl Te cells. Data represent one of three independent experiments. (**B-E**) An enhanced autoimmune response in Nrn1^-/-^ mice *in vivo*. (**B**) Experimental scheme. Nrn1^-/-^ mice were crossed with FDG mice and Nrn1^-/-^_FDG or ctrl_FDG mice were obtained. The autoimmune response was induced by injecting DT i.p. to delete endogenous Treg cells. Mice’s weight change was monitored after disease induction. (**C**) Relative body weight change after autoimmune response induction. (**D**) Mice were harvested six days after DT injection and assessed for ki67, cytokine TNFα, IL2, and IFNγ expression in CD4^+^ cells. (**E**) Foxp3 expression among CD4^+^ cells day six post DT treatment. n≥5 mice per group. Data represent four independent experiments. (**F-I**) Changes relating to ion balances in Te cells. (**F**) Gene sets clusters from GSEA of GO_MF and GO_Biological process (GO_BP) results in Nrn1^-/-^ and ctrl Te cells (Figure 4-figure supplement Table 4). (**G**) Enrichment of “GOBP_ membrane repolarization” gene set and enriched gene expression heatmap. (**H**) Membrane potential measurement in Te cells. Data represent two independent experiments. (**I**) Enrichment of “GOMF_Metal ion transmembrane transporter activity” gene set and heatmap of differential gene expression pattern (Figure 4-figure supplement 1B). (**J-N**) Metabolic changes associated with Nrn1^-/-^ Te cell. (**J**) Enrichment of “GOMF_amino acid transmembrane transporter activity” gene set and differential gene expression heatmap. (**K**) AAs induced MP changes in Te cells. Data represent two independent experiments. (**L**) Measurement of pmTOR and pS6 in Te cells after nutrient sensing. Data represent three independent experiments. (**M**) Enriched Hallmark gene sets (p<0.05, FDR q<0.25). (**N**) Seahorse analysis of extracellular acidification rate (ECAR) and oxygen consumption rate (OCR) in Nrn1^-/-^ and ctrl Te cells. n≥6 technical replicates per group. Data represent three independent experiments. Error bars indicate ±SEM. *p<0.05, **p<0.01, ***p<0.001, ****p<0.0001, unpaired Student’s t-test was performed for two-group comparison.

To identify molecular changes responsible for Nrn1^-/-^ Te phenotype, we compared gene expression between Nrn1^-/-^ and ctrl Te cells by RNASeq. GSEA and Cytoscape analysis identified a cluster of gene sets on “membrane repolarization”, suggesting that Nrn1 may also be involved in the regulation of MP under Te context (Figure 4F, Figure 4-figure supplement Table 4) (Shannon et al., 2003; Subramanian et al., 2005). While the “membrane_repolarization” gene set was enriched in Nrn1^-/-^ (Figure 4G), the “Neurotransmitter receptor activity involved in regulation of postsynaptic membrane potential” gene set was no longer enriched, but the AMPAR subunit Gria3 expression was still elevated in Nrn1^-/-^ Te cells (Figure 4-figure supplement 1A). Although MP in Te cells was comparable between Nrn1^-/-^ and ctrl (Figure 4H), the “MF_metal ion transmembrane transporter activity” gene set was significantly enriched in Nrn1^-/-^ with different gene expression patterns (Figure 4I, Figure 4-figure supplement 1B), indicative of different electric state. The significant enrichment of ion channel related genes in Nrn1^-/-^ Te cells was in line with the finding in Nrn1^-/-^ iTreg cells, supporting the notion that Nrn1 expression may be involved in ion balance and MP modulation.

Examination of nutrient transporters revealed that the “Amino acid transmembrane transporter activity” gene set was significantly enriched in Nrn1^-/-^ cells than the ctrl (Figure 4J). We further examined AA entry-induced cellular MP change in Nrn1^-/-^ and ctrl Te cells. AA entry caused enhanced MP change among Nrn1^-/-^ Te than the ctrl, in contrast with the finding under iTreg cell context (Figure 4K). Along with the enrichment of ion channel and nutrient transporter genes (Figure 4I and J), we found enhanced mTOR and S6 phosphorylation in Nrn1^-/-^ Te cells (Figure 4-supplemental figure 1C). We also compared nutrient sensing capability between Nrn1^-/-^ and ctrl Te cells, as outlined in Figure 3I. Nrn1^-/-^ Te showed increased mTOR and S6 phosphorylation after sensing ions and nutrients in RPMI medium (Figure 4L), confirming the differential impact of ions and nutrients on Nrn1^-/-^ and ctrl Te cells. GSEA on Hallmark collection showed enrichment of mTORC1 signaling gene set (Figure 4M), corroborating with increased pmTOR and pS6 detection in Nrn1^-/-^ Te cells. Along with increased mTORC1 signaling, Nrn1^-/-^ Te cells also showed enrichment of gene sets on glycolysis and proliferation (Figure 4M). Evaluation of metabolic changes by seahorse confirmed increased glycolysis in Nrn1^-/-^ cells, while the OCR remained comparable between Nrn1^-/-^ and ctrl (Figure 4N). These *in vitro* studies on Te cells indicate that Nrn1 deficiency resulted in the dysregulation of the electrolyte and nutrient transport program, impacting Te cell nutrient sensing, metabolic state, and the outcome of inflammatory response.

### Nrn1 deficiency exacerbates autoimmune disease

The coordinated reaction of Treg and Te cells contributes to the outcome of the immune response. We employed the experimental autoimmune encephalomyelitis (EAE), the murine model of multiple sclerosis (MS) to evaluate the overall impact of Nrn1 on autoimmune disease development. Upon EAE induction, the incidence and time to EAE onset in Nrn1^-/-^ mice were comparable to the ctrl mice, but the severity, disease persistence, and body weight loss were increased in Nrn1^-/-^ mice (Figure 5A). Exacerbated EAE was associated with significantly increased CD45^+^ cell infiltrates, increased CD4^+^ cell number, increased proportion of MOG-specific CD4 cells, and reduced proportion of Foxp3^+^ CD4 cells in the Nrn1^-/-^ spinal cord (Figure 5B-E). Moreover, we also observed increased proportions of IFNγ^+^ and IL17^+^ CD4 cells in Nrn1^-/-^ mice remaining in the draining lymph node compared to the ctrl mice (Figure 5F). Thus, the results from EAE corroborated with earlier data and confirmed the important role of Nrn1 in establishing immune tolerance and modulating autoimmunity.

**Figure 5.**
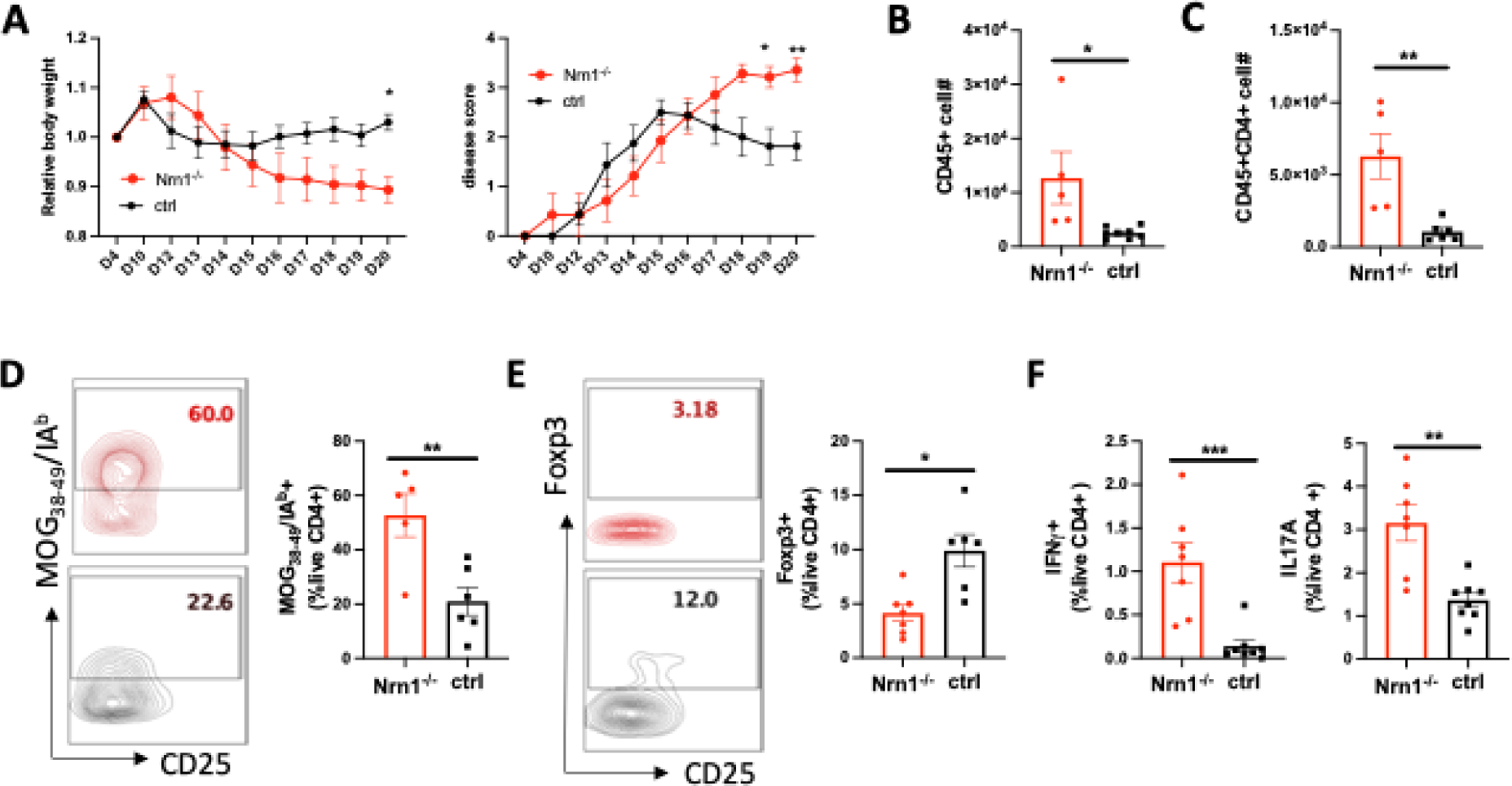
Nrn1 deficiency exacerbates autoimmune EAE disease. (**A**) Aggravated body weight loss and protracted EAE disease in Nrn1^-/-^ mice. (**B**) CD45^+^ cell number in the spinal cord infiltrates. (**C**) CD4^+^ cell number in the spinal cord infiltrates. (**D**) Mog_38-49_/IA^b^ tetramer staining of spinal cord infiltrating CD4 cells. (**E**) Foxp3^+^ proportion among CD4^+^ cells in spinal cord infiltrates. (**F**) IFNγ^+^ and IL17^+^ cell proportion among CD4^+^ cells in draining lymph nodes. n≥5 mice per group. Data represent three independent experiments. The P value was calculated by 2way ANOVA for (**A**). The p-value was calculated by the unpaired student t-test for (**B-F**). \**P*<0.05, \*\**P*<0.01.

## Discussion

T cell expansion and functional development depend on adaptive electric and metabolic changes, maintaining electrolyte balances, and appropriate nutrient uptake. The negative charge of the plasma membrane, ion channel expression pattern, and function are key characteristics associated with the cellular electric state in different systems, impacting cell proliferation and function (Blackiston et al., 2009; Emmons-Bell and Hariharan, 2021; Kiefer et al., 1980; Monroe and Cambier, 1983; Sundelacruz et al., 2009). The electrolytes and nutrients, including amino acids, metabolites, and small peptides transported through ion channels and nutrient transporters, are also regulators and signaling agents impacting the choice of cellular metabolic pathways and functional outcomes (Hamill et al., 2020). In this study, we report that the neurotropic factor Nrn1 expression influences CD4 T cell MP, ion channels, and nutrient transporter expression patterns, contributing to differential metabolic states in Treg and Te cells. Nrn1 deficiency compromises Treg cell expansion and suppression while enhancing Te cell inflammatory response, exacerbating autoimmune disease.

Bioelectric controls have been defined as a type of epigenetics that can control information residing outside of genomic sequence (Levin, 2021). The sum of ion channels and pump activity generates the ionic gradient across the cell membrane, establishing the MP level and bioelectric state. Cells with the same MP can have different ion compositions, and the same ion channel may have a differential impact on MP when in combination with different ion channels (Abdul Kadir et al., 2018). Consistent with this notion, Nrn1 deficiency has differential impacts on the cellular electric state under the Treg and Te cells with different ion channel combinations. Altered MP was detected in Nrn1 deficient Treg cells (Figure 3E), while comparable MP was observed between Nrn1^-/-^ and ctrl Te cells (Figure 4H). The MP level determined by ion channels and pump activity can influence the nutrient transport pattern, establishing a metabolic and functional state matching the MP level (Blackiston et al., 2009; Emmons-Bell and Hariharan, 2021; Kiefer et al., 1980; Monroe and Cambier, 1983; Sundelacruz et al., 2009; Yu et al., 2022). Yu et al. reported that macrophage MP modulates plasma membrane phospholipid dynamics and facilitates cell surface retention of nutrient transporters, thus supporting nutrient uptake and impacting the inflammatory response (Yu et al., 2022). Nutrient transport is key to T cell fate decisions and has been considered signal 4 to T cell fate choices (Chapman and Chi, 2022; Long et al., 2021). The changes in ion channel related gene expression and MP level in Nrn1^-/-^ cells were accompanied by differential expression of AA transporter genes and nutrient sensing activity that impacted mTORC1 pathway activation and cellular glycolytic state (Figures 3 and 4). These results corroborate previous observations on the connection of MP in nutrient acquisition and metabolic change and support the role of Nrn1 in coordinating T cell electric and metabolic adaptation (Yu et al., 2022).

Although Nrn1, as a small GPI-anchored protein, does not have channel activity by itself, it has been identified as one of the components in the AMPAR complex (Pandya et al., 2018; Schwenk et al., 2012; Subramanian et al., 2019). Na^+^-influx through the AMPA type ionotropic glutamate receptor can quickly depolarize the postsynaptic compartment and potentiate synaptic transmission in neurons. We have observed increased expression of AMPAR subunits in Nrn1^-/-^ iTreg and Te cells (Figure 3D, Figure 4-figure supplement 1), implicating potential change in AMPAR activity in Nrn1^-/-^ under Treg and Te cell context. Glutamate secreted by proliferating cells may influence T cell function through AMPAR. High glutamate levels are detected at the autoimmune disease site and tumor interstitial fluid (Bonnet et al., 2020; McNearney et al., 2004; Sullivan et al., 2019). Moreover, AMPAR has been implicated in exacerbating autoimmune disease (Bonnet et al., 2015; Sarchielli et al., 2007). The increased expression of AMPAR subunits in Nrn1^-/-^ cells supports the potential connection of Nrn1 and AMPAR and warrants future investigation on the possibility that Nrn1 functions through AMPAR, impacting T cell electric change. Besides AMPAR, Nrn1 has been reported to function through the insulin receptor and fibroblast growth factor pathway (Shimada et al., 2016; Yao et al., 2012). Subramanian et al have suggested that rather than a traditional ligand with its cognate receptor, Nrn1 may function as an adaptor to receptors to perform diverse cell-type-specific functions (Subramanian et al., 2019). Our results do not rule out these possibilities.

Overall, we found that Nrn1 expression in Treg and Te cells can impact cellular electric state, nutrient sensing, and metabolism in a cell context dependent manner. The predominant enrichment of ion channel related gene sets in both Treg and Te cell context underscores the importance of Nrn1 in modulating ion balance and MP. The changes in ion channels and nutrient transporter expression in Treg and Te cells and associated functional consequences highlight the importance of Nrn1 in coordinating cell metabolic changes through channels and transporters during the adaptive response and contribute to the balance of tolerance and immunity.

## Materials and Methods

### Mouse models

The Nrn1^-/-^ mice (Fujino et al., 2011), Foxp3DTRGFP (FDG)(Kim et al., 2007), and TCRα^-/-^ mice were obtained from the Jackson Laboratory. OTII mice on Thy1.1^+^ background was kindly provided by Dr. Jonathan Powell. Rag2^-/-^ mice were maintained in our mouse facility. 6.5 TCR transgenic mice specific for HA antigen and C3HA mice (both on the B10.D2 background) have been described previously (Huang et al., 2004). Nrn1^-/-^ mice were crossed with OTII mice to generate Nrn1^-/-^_OTII^+^ mice, ctrl_OTII^+^ mice. Nrn1^-/-^ mice were also crossed with FDG mice to generate Nrn1^-/-^_FDG and ctrl_FDG mice. All mice colonies were maintained in accordance with the guidelines of Johns Hopkins University and the institutional animal care and use committee

### Antibodies and Reagents

We have used the following antibodies: Anti-CD3 (17A2), anti-CD4 (RM4-5), anti-CD8a (53-6.7), anti-CD25 (PC61), anti-CD45.1 (A20), anti-CD45.2 (104), anti-CD62L (MEL-14), CD73 (TY/111.8), anti-CD90.1 (OX-7), anti-CD90.2 (30-H12), anti-TCR V 5.1, 5.2 (MR9-4), anti- PD1 (29F.1A12), anti-IFNγ (XMG1.2), anti-IL17α (TC11-18H10.1), anti-TNFα (MP6-XT22), anti-Tbet (4B10), anti-Ki67 (16A8) were purchased from Biolegend. Anti-CD44 (IM7), CD45 (30-F11), anti-CD69(H1.2F3) were purchased from BD Bioscience. Anti-FOXP3 (FJK-16s) was purchased from eBioscience. The flow cytometry data were collected using BD Celesta (BD Biosciences) or Attune Flow Cytometers (ThermoFisher). Data were analyzed using FlowJo (Tree Star) software.

Mouse monoclonal anti-Nrn1 antibody (Ab) against Nrn1 was custom-made (A&G Pharmaceutical). The specificity of anti-Nrn1 Ab was confirmed by ELISA, cell surface staining of Nrn1 transfected 293T cells, and western blot of Nrn1 recombinant protein and brain protein lysate from WT mice or Nrn1^-/-^ mice (data not shown). OVA_323-339_ peptide and MOG_35-55_ was purchased from GeneScript. Incomplete Freund’s adjuvant (IFA) and Mycobacterium tuberculosis H37Ra (killed and desiccated) were purchased from Difco. Pertussis toxin was purchased from List Biological Laboratories and diphtheria toxin was obtained from Millipore-Sigma.

### Cell purification and culture

Naïve CD4 cells were isolated from the spleen and peripheral lymph node by a magnetic bead-based purification according to the manufacturer’s instruction (Miltenyi Biotech). Purified CD4 cells were stimulated with plate-bound anti-CD3 (5ug/ml, Bio-X-Cell) and anti-CD28 (2ug/ml, Bio-X-Cell) for 3 days, in RPMI1640 medium supplemented with 10%FBS, HEPES, penicillin/streptomycin, MEM Non-Essential Amino Acids, and β-mercaptoethanol. For iTreg cell differentiation, cells were stimulated in the presence of human IL2 (100u/ml, PeproTech), human TGFβ (10ng/ml, PeproTech), anti-IL4, and anti-IFNγ antibody (5ug/ml, Clone 11B11 and clone XMG1.2, Bio-X-Cell) in 10% RPMI medium. CD4^+^ Te cells were differentiated without additional cytokine or antibody for three days, followed by additional culture for 2 days in IL2 100u/ml in 10%RPMI medium. nTreg cells were isolated by sorting from the FDG CD4^+^ fraction based on Foxp3^+^GFP and CD25 expression (CD4^+^CD25^+^GFP^+^). Alternatively, nTreg cells were enriched from CD4 cells by positive selection using the CD4^+^CD25^+^ Regulatory T Cell Isolation Kit from Miltenyi.

### Self-antigen induced tolerance model

1x10^6^ HA-specific Thy1.1^+^ 6.5 CD4 cells from donor mice on a B10.D2 background were transferred into C3-HA recipient mice, where HA is expressed as self-antigen in the lung; or into WT B10.D2 mice followed by infection with Vac-HA virus (1x10^6^ pfu). HA-reactive T cells were recovered from the lung-draining lymph node of C3-HA host mice or WT B10.D2 Vac-HA infected mice at indicated time points by cell sorting. RNA from sorted cells was used for qRT-PCR assay examining Nrn1 expression.

### Peptide-induced T cell anergy model

5x10^5^ Polyclonal Treg cells from CD45.1^+^ C57BL/6 mice were mixed with 5x10^6^ thy1.1^+^ OTII cells from Nrn1^-/-^_OTII or ctrl_OTII mice and transferred by *i.v.* injection into TCRα^-/-^ mice. 100ug of OVA_323–339_ dissolved in PBS was administered *i.v.* on days 1, 4, and 7 after cell transfer. Host mice were harvested on day 13 after cell transfer, and cells from the lymph node and spleen were further analyzed.

### *In vivo* Treg suppression assay

nTreg cells from CD45.2^+^CD45.1^-^ Nrn1^-/-^ or ctrl mice (5x10^5^/mouse) in conjunction with CD45.1^+^ splenocytes (2x10^6^/mouse) from FDG mice were cotransferred i.p. into Rag2^-/-^ mice. The CD45.1^+^ splenocytes were obtained from FDG mice pretreated with DT for 2 days to deplete Treg cells. Treg suppression toward CD45.1^+^ responder cells was assessed on day 7 post cell transfer. Alternatively, 7 days after cell transfer, Rag2^-/-^ hosts were challenged with an *i.d.* inoculation of B16F10 cells (1x10^5^). Tumor growth was monitored daily. Treg-mediated suppression toward anti-tumor response was assessed by harvesting mice day 18-21 post-tumor inoculation.

### Induction of autoimmunity by transient Treg depletion

To induce autoimmunity in Nrn1^-/-^_FDG and ctrl_FDG mice, 1ug/mouse of DT was administered i.p. for two consecutive days, and the weight loss of treated mice was observed over time.

### EAE induction

EAE was induced in mice by subcutaneous injection of 200 μg MOG_35–55_ peptide with 500 μg M. tuberculosis strain H37Ra (Difco) emulsified in incomplete Freund Adjuvant oil in 200ul volume into the flanks at two different sites. In addition, the mice received 400 ng pertussis toxin (PTX; List Biological Laboratories) *i.p*. at the time of immunization and 48 h later. Clinical signs of EAE were assessed daily according to the standard 5-point scale (Miller et al., 2007): normal mouse; 0, no overt signs of disease; 1, limp tail; 2, limp tail plus hindlimb weakness; 3, total hindlimb paralysis; 4, hindlimb paralysis plus 75% of body paralysis (forelimb paralysis/weakness); 5, moribund.

### ELISA

MaxiSorp ELISA plates (ThermoScientific Nunc) were coated with 100 μl of 1μg/ml anti-mIL-2 (BD Pharmingen #554424) at 4°C overnight. Coated plates were blocked with 200μl of blocking solution (10%FBS in PBS) for 1hr at room temperature (RT) followed by incubation of culture supernatant and mIL-2 at different concentrations as standard. After 1hr, plates were washed and incubated with anti-mIL-2-biotin (BD Pharmingen #554426) at RT for 1hr. After 1hr, plates were incubated with 100μl of horseradish peroxidase-labeled avidin (Vector Laboratory, #A-2004) 1μg/ml for 30min. After washing, samples were developed using the KPL TMB Peroxidase substrate system (Seracare #5120-0047) and read at 405 or 450 nm after the addition of the stop solution.

### Quantitative RT-PCR

RNA was isolated using the RNeasy Micro Kit (Qiagen 70004) following the manufacturer’s instructions. RNA was converted to cDNA using the High-Capacity cDNA Reverse Transcription Kit (ThermoFisher Scientific #4368814) according to the manufacturer’s instructions. The primers of murine genes were purchased from Integrated DNA Technology (IDT). qPCR was performed using the PowerUp SYBR Green Master Mix (ThermoFisher Scientific #A25780) and the Applied Biosystems StepOnePlus 96-well real-time PCR system. Gene expression levels were calculated based on the Delta-Delta Ct relative quantification method. Primers used for Nrn1 PCR were as follows: GCGGTGCAAATAGCTTACCTG (forward); CGGTCTTGATGTTCGTCTTGTC (reverse).

### Ca^++^ flux and Membrane potential measurement

To measure Ca^++^ flux, CD4 cells were loaded with Fluo4 dye at 2μM in the complete cell culture medium at 37°C for 30min. Cells were washed and resuspended in HBSS Ca^++^ free medium and plated into 384 well glass bottom assay plate (minimum of 4 wells per sample). Ca^++^ flux was measured using the FDSS6000 system (Hamamatsu Photonics). To measure store-operated calcium entry (SOCE), after the recording of the baseline T cells Ca^++^ fluorescent for 1min, thapsigargin (TG) was added to induce store Ca^++^ depletion, followed by the addition of Ca^++^ 2μM in the extracellular medium to observe Ca^++^ cellular entry.

Membrane potential was measured using FLIPR Membrane Potential Assay kit (Molecular devices) according to the manufacturer’s instructions. Specifically, T cells were loaded with FLIPR dye by adding an equal volume of FLIPR dye to the cells and incubated at 37°C for 30 minutes. Relative membrane potential was measured by detecting FLIPR dye incorporation using flow cytometry.

To measure changes of MP after AAs transport, T cells were plated and loaded with FLIPR dye at 37°C for 30 minutes in 384 well glass bottom assay plate (minimum of 6 wells per sample). After recording the baseline T cell MP for 1min, MEM AAs (Gibco MEM Amino Acids #11130-051) were injected into each well, and the change of MP was recorded for 5 min.

### Extracellular flux analysis (Seahorse assays)

Real-time measurements of extracellular acidification rate (ECAR) and oxygen consumption rate (OCR) were performed using an XFe-96 Bioanalyser (Agilent). T cells (2 × 10^5^ cells per well; minimum of four wells per sample) were spun into previously poly-d-lysine-coated 96-well plates (Seahorse) in complete RPMI-1640 medium. ECAR was measured in RPMI medium in basal condition and in response to 25mM glucose, 1μM oligomycin, and 50mM of 2-DG (all from Sigma Aldrich). OCR was measured in RPMI medium supplemented with 25mM glucose, 2mM L-glutamine, and 1mM sodium pyruvate, under basal condition and in response to 1μM oligomycin, 1.5μM of carbonylcyanide-4-(trifluoromethoxy)-phenylhydrazone (FCCP) and 1μM of rotenone and antimycin (all from Sigma Aldrich).

### RNAseq and data analysis

RNASeq samples: 1. Anergic T cell analysis. Ctrl and Nrn1^-/-^ OTII cells were sorted from the host mice (n=3 per group). 2. iTreg cell analysis. In vitro differentiated Nrn1^-/-^ and ctrl iTreg cells were replated in resting condition (IL2 100u/ml) or stimulation condition (IL2 100u/ml and aCD3 5ug/ml). Cells were harvested 20 hr after replating for RNASeq analysis. 3. Effector T cells. Nrn1^-/-^ and ctrl CD4 Tn cells were activated for 3 days (aCD3 5ug/ml, aCD28 2ug/ml), followed by replating in IL2 medium (100u/ml). Te cells were harvested two days after replating and subjected to RNASeq analysis.

RNA-sequencing analysis was performed by Admera Health (South Plainfield, NJ). Read quality was assessed with FastQC and aligned to the Mus Musculus genome (Ensembl GRCm38) using STAR aligner (version 2.6.0)(Dobin et al., 2013). Aligned reads were counted using HTSeq (version 0.9.0)(Anders et al., 2015), and the counts were loaded into R (The R Foundation). DESeq2 package (version 1.24.0)(Love et al., 2014) was used to normalize the raw counts. GSEA was performed using public gene sets (HALLMARK, and GO)(Subramanian et al., 2005). Cytoscape was used to display enriched gene sets cluster (Shannon et al., 2003).

Statistical analysis. All numerical data were processed using Graph Pad Prism 10. Data are expressed as the mean +/- the SEM, or as stated. Statistical comparisons were made using an unpaired student t-test or ANOVA with multiple comparison tests where 0.05 was considered significant, and a normal distribution was assumed. The p values are represented as follows: * p<0.05; ** p<0.01; *** p<0.001, **** p<0.0001.

## Acknowledgements

This research is supported by grants from the Bloomberg-Kimmel Institute of JHU, the Melanoma Research Alliance, the National Institutes of Health (RO1AI099300 and RO1AI089830), and the Department of Defense (PC130767). JB’s research was supported by a Crohn’s and Colitis Foundation of America Research Fellowship, the Melanoma Research Foundation, and NCI grant P30CA016056.

We thank Dennis Gong for data processing and critical reading of the manuscript. We thank Dr. Elly Nedivi for providing polyclonal Nrn1 antibody and Dr. Fan Pan for reagent support. We thank Drs. Franck Housseau, Chien-Fu Hung for the constructive discussion of the project and the manuscript. We thank Drs. Hao Shi and Hongbo Chi for critical reading of the manuscript and helpful suggestions. We thank Dr. Rachel Helm for manuscript editing. We thank Drs. Richard L. Huganir, Bian Liu and Hana Goldschmidt for constructive discussion on Nrn1 and AMPAR connection.

## Author contributions

H. Y. was involved in all aspects of this study, including planning and performing experiments, analyzing and interpreting data, and writing the manuscript. H.N. and J.B. were involved in performing experiments and data interpretation. P.V., Y.Z, and A.L. analyzed Nrn1 expression in Treg cells and performed Treg suppression and functional assay. M.M. was involved in Nrn1 expression, Treg suppression assay, and manuscript writing. C. H. and C.D. conducted the anergy and Te cell differential gene expression study; Y.Z., J.F., and K.C. conducted an autoimmune inflammation study, and M.H. helped with mouse colony genotyping. X.Z. and Z.L. contributed to bioinformatic analysis. D.M.P. oversaw the project and was involved in data interpretation, manuscript preparation, and funding support.

## Competing interests

C.D. is a co-inventor on patents licensed from JHU to BMS and Janssen and is currently an employee of Janssen Research. D.M.P. is a consultant for Compugen, Shattuck Labs, WindMIL, Tempest, Immunai, Bristol-Myers Squibb, Amgen, Janssen, Astellas, Rockspring Capital, Immunomic, and Dracen; owns founders equity in ManaT Bio Inc., WindMIL, Trex, Jounce, Enara, Tizona, Tieza, and RAPT; and receives research funding from Compugen, Bristol-Myers Squibb, and Enara. All other authors do not have conflicting financial interests.

## Data availability

The raw RNA sequencing data has been deposited under the GEO accession no. GSE121908 and GSE224083.

## Figure supplements

**Figure 1-figure supplement 1.**
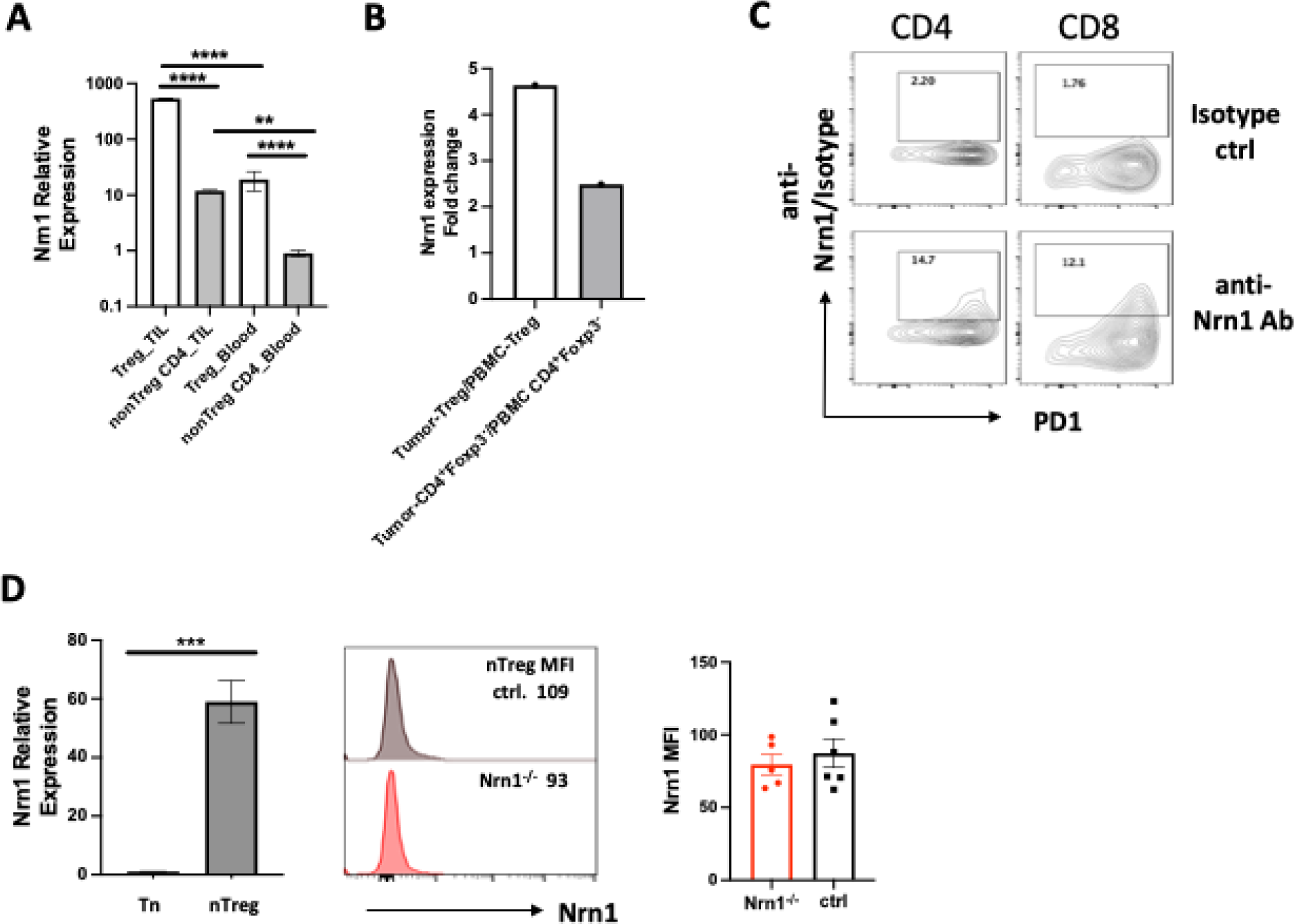
Nrn1 expression in T cells from tumor environment and during early T cell activation. **(A-B).** Nrn1 expression in tumor infiltrates. (A) Comparison of Nrn1 expression by qRT-PCR among Tregs and non-Treg CD44^hi^CD4^+^ cells recovered either from B16 melanoma infiltrates or from peripheral blood of Foxp3DTRgfp mice bearing subcutaneous B16 melanomas. **(B)** Comparison of Nrn1 expression in breast tumor-infiltrating Treg (T-Treg) and Te (T-CD4^+^Foxp3^-^) cells vs. peripheral blood Treg (P-Treg) and Te (PBMC CD4^+^Foxp3^-^) cells. Data derived from the “Regulatory T Cells Exhibit Distinct Features in Human Breast Cancer” report (Plitas et al., 2016). **(C)** Nrn1 cell surface detection on day 2 activated CD4 and CD8 cells by flow cytometry. **(D).** Detection of Nrn1 expression in nTreg cells by qRT-PCR and cell surface Nrn1 staining.

**Figure 1-figure supplement 2.**
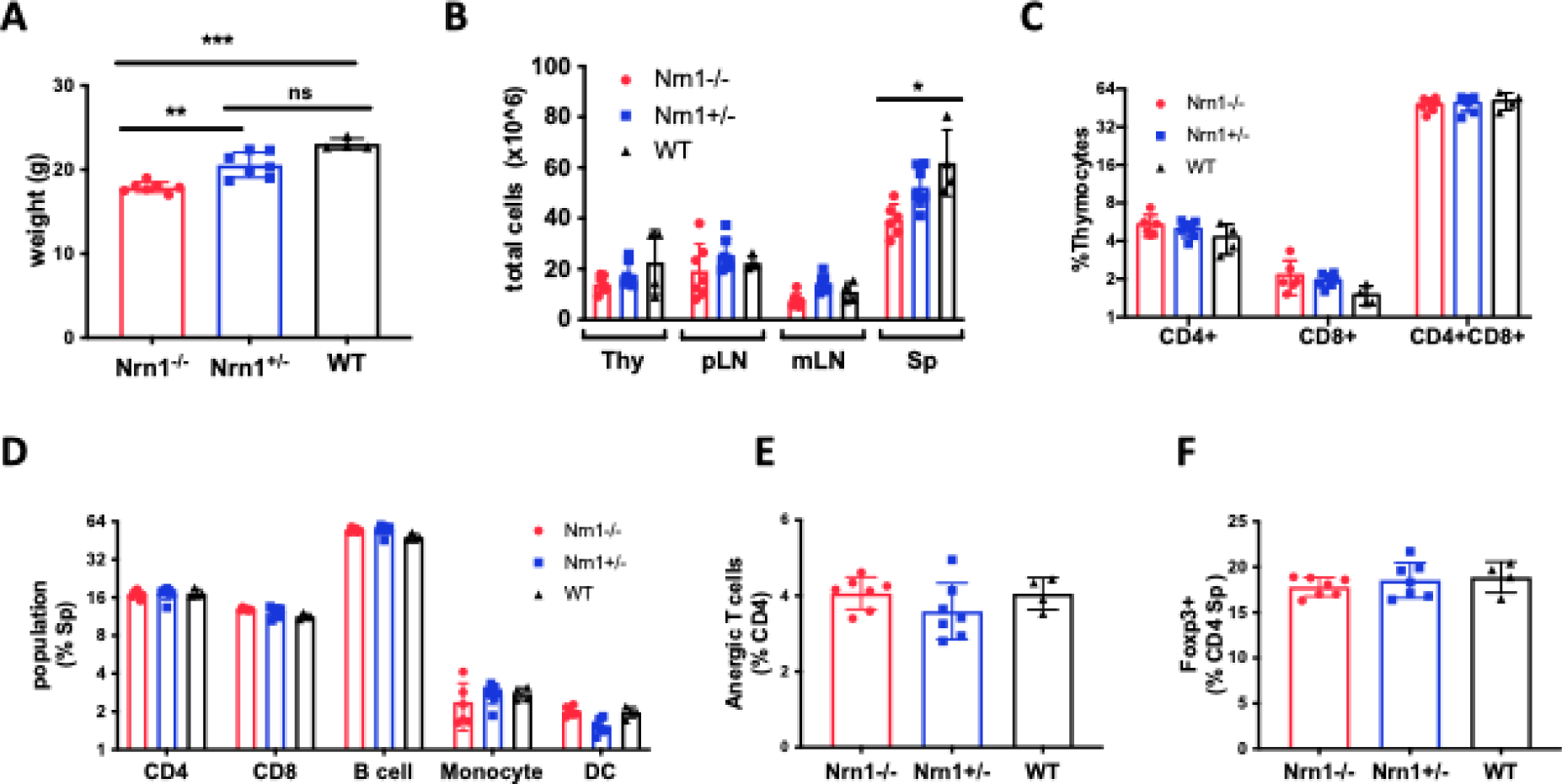
Nrn1^-/-^ mice body weight and immune cell profile analysis compared to Nrn1^+/-^ and WT mice. **(A)** Average body weight of 10-12 wk old age and sex-matched Nrn1^-/-^, Nrn1^+/-^ and WT mice. **(B)** Thymus and peripheral lymphoid tissue total cell count. **(C-D)** Immune cell frequencies in the thymus and spleen. **(E)** Proportion of CD4^+^Foxp3^-^ CD44^+^FR4^hi^ CD73^hi^ anergic T cells among splenocytes CD4 cell population. **(F)** FOXP3^+^cell frequency among CD4 cells in the spleen. The immune profile assessment used n>3 mice/group of Nrn1^-/-^ and control mice. *P* values were calculated by one-way analysis of variance (ANOVA). \**P*<0.05. \*\**P*<0.01.

**Figure 1-figure supplement 3.**
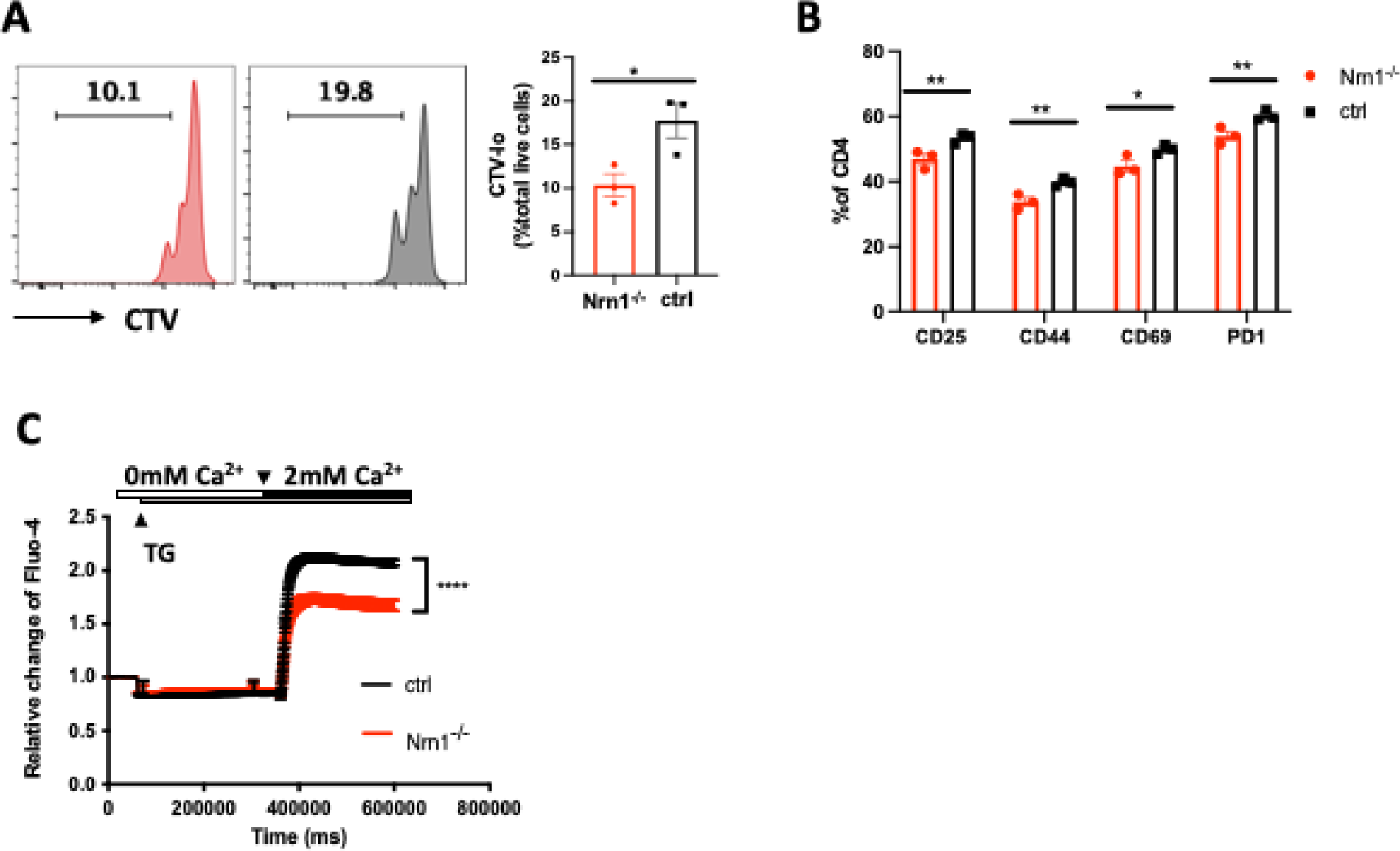
Compromised T cell activation in Nrn1^-/-^ cells. **(A)** Cell tracker dye violet (CTV) dilution in Nrn1^-/-^ or ctrl CD4 cells after stimulation with plate-bound aCD3 (5ug/ml) and soluble aCD28 (2ug/ml); **(B)** Cell surface activation markers CD25, CD44, CD69, and PD1 expression day 2 after naïve CD4^+^ cell activation. Unpaired student’s t-test, *p<0.05, **p<0.01. Data represent three independent experiments. **(C)** Store-operated Ca^++^ entry (SOCE) was examined on day 2 activated CD4^+^ cells labeled with Fluo-4 dye. Representative graph and mean ± SEM of SOCE induced by CD4 cell stimulation with 1uM thapsigargin (TG) in Ca^++^ free HBSS (0mM Ca^++^) followed by addition of 2mM Ca^++^. Representative graphs of Ca^++^ influx from three independent experiments (****p<0.0001).

**Figure 3-figure supplement 1.**
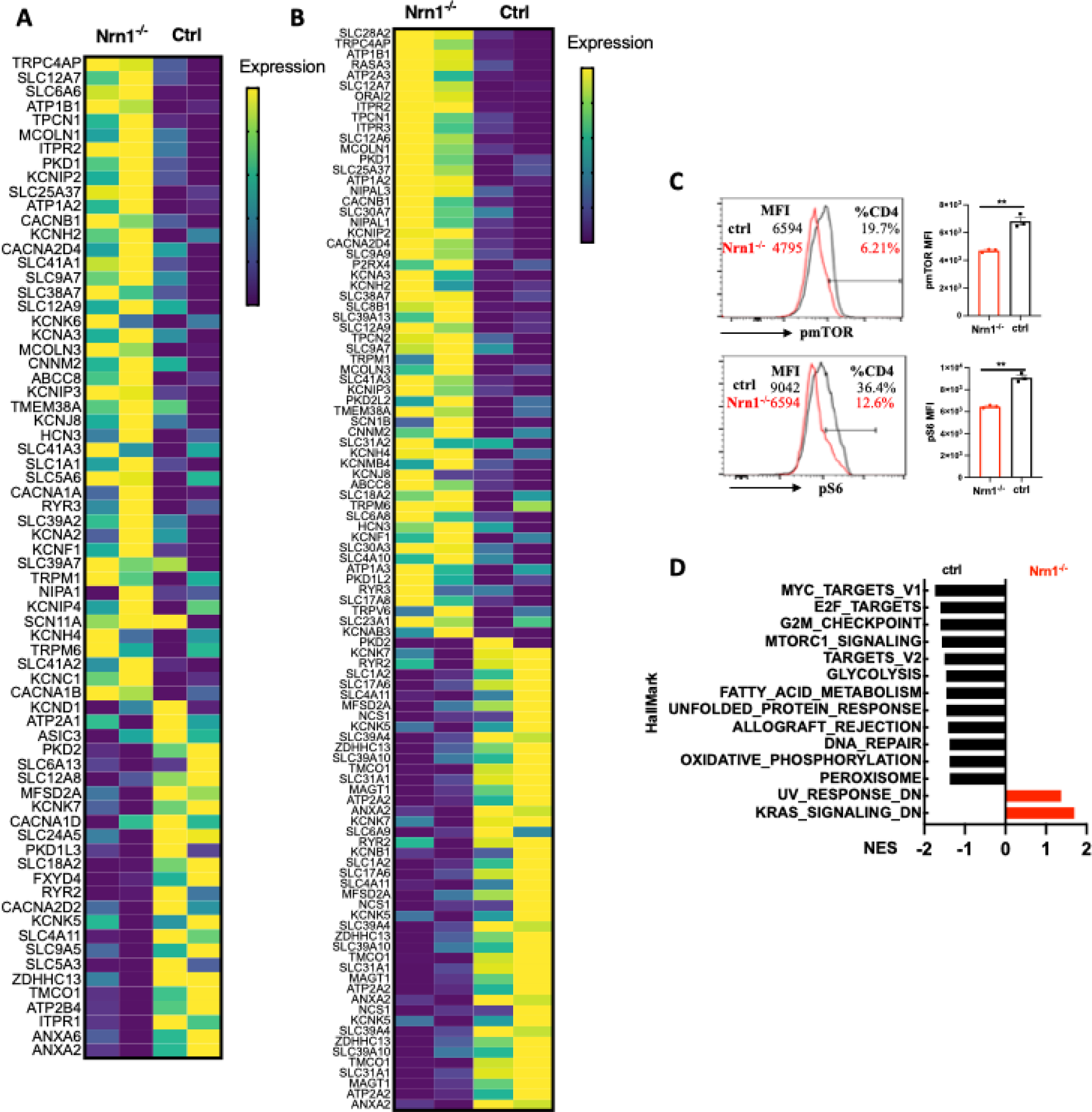
Heatmap of differentially expressed genes and Hallmark gene set enrichment. **(A)** Heatmap of differentially expressed genes in “GOMF_Metal ion transmembrane transporter activity” gene set from Nrn1^-/-^ and ctrl iTreg cells cultured under the resting condition. **(B)** Heatmap of differentially expressed genes in “GOMF_Metal ion transmembrane transporter activity” gene set from reactivated Nrn1^-/-^ and ctrl iTreg cells. **(C)** Detection of pmTOR and pS6 in Nrn1^-/-^ and ctrl iTreg cells. Data represents three independent experiments. **p<0.01. Unpaired Student’s t-tests were performed. **(D).** Enrichment of Hallmark gene set in activated Nrn1^-/-^ and ctrl iTreg cells (P<0.05, FDR q<0.25).

**Figure 3-figure supplement 2.**
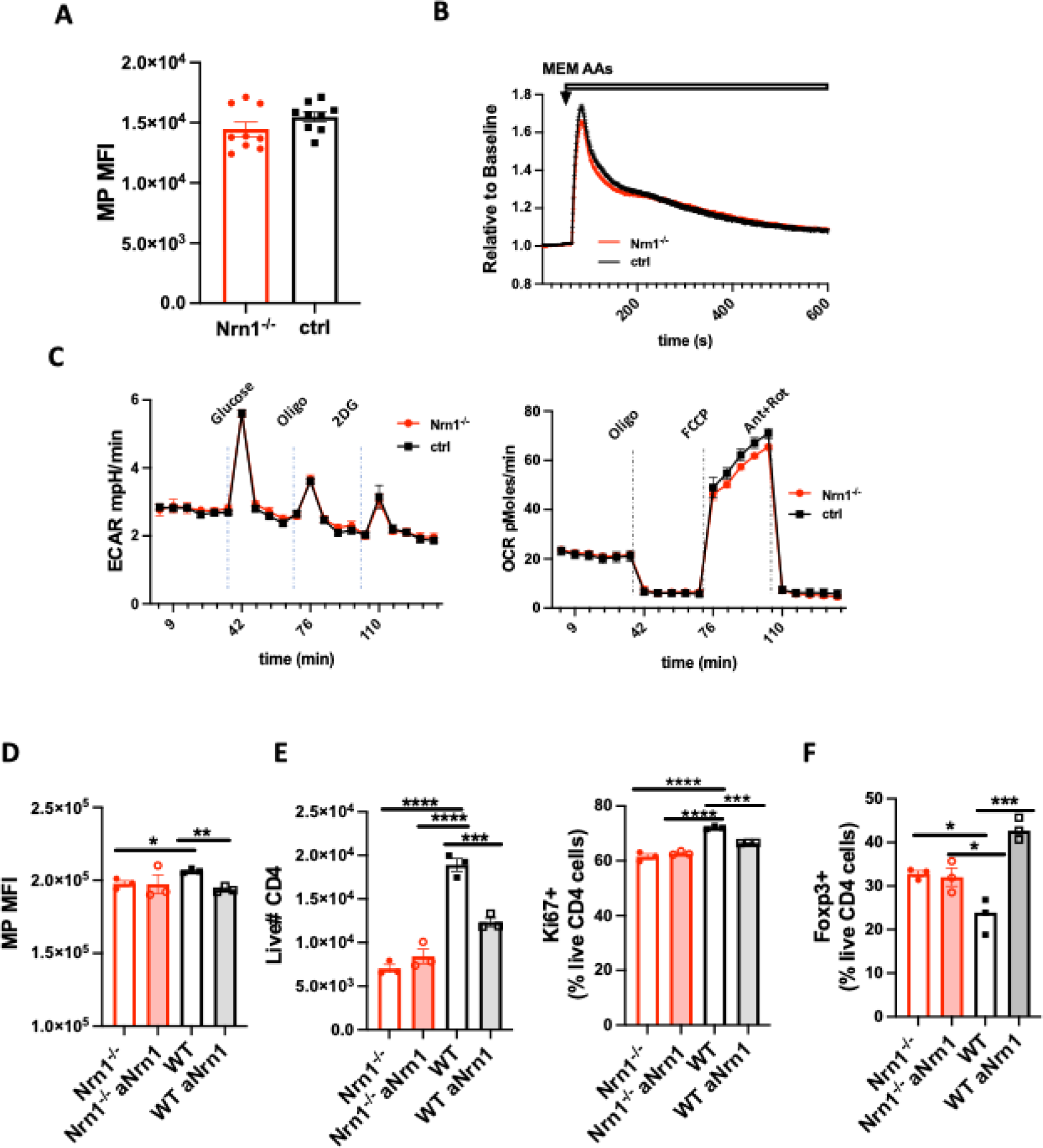
Characterization of Nrn1^-/-^ naïve CD4 T cells and effect of Nrn1 blockade on WT iTreg cell differentiation and expansion. **(A)** Resting MP in Nrn1^-/-^ and ctrl naïve CD4^+^ T cells. **(B)** AAs induced MP change in Nrn1^-/-^ and ctrl naïve CD4^+^ T cells. (**C)** Seahorse analysis of extracellular acidification rate (ECAR) and oxygen consumption rate (OCR) in Nrn1^-/-^ and ctrl iTreg cells. n=6∼10 technical replicates per group. Data represent three independent experiments. **(D-F)** WT iTreg cells differentiated in the presence of Nrn1 antibody blockade. **(D).** MP in Nrn1^-/-^ and WT iTreg cells differentiated in the presence or absence of anti-Nrn1 antibody. **(E).** Live cell number and proportion of Ki67 expressing cells after anti-CD3 restimulation among Nrn1^-/-^ and WT iTreg cells differentiated in the presence or absence of anti-Nrn1 antibody. **(F).** Foxp3^+^ cell proportion three days after anti-CD3 restimulation in Nrn1^-/-^ and WT iTreg cells differentiated and restimulated in the presence or absence of anti-Nrn1 antibody. Data represent three independent experiments. *p<0.05, **p<0.01, ***p<0.001, ****p<0.0001. Ordinary one-way ANOVA was performed for multi-comparison.

**Figure 4-figure supplement 1.**
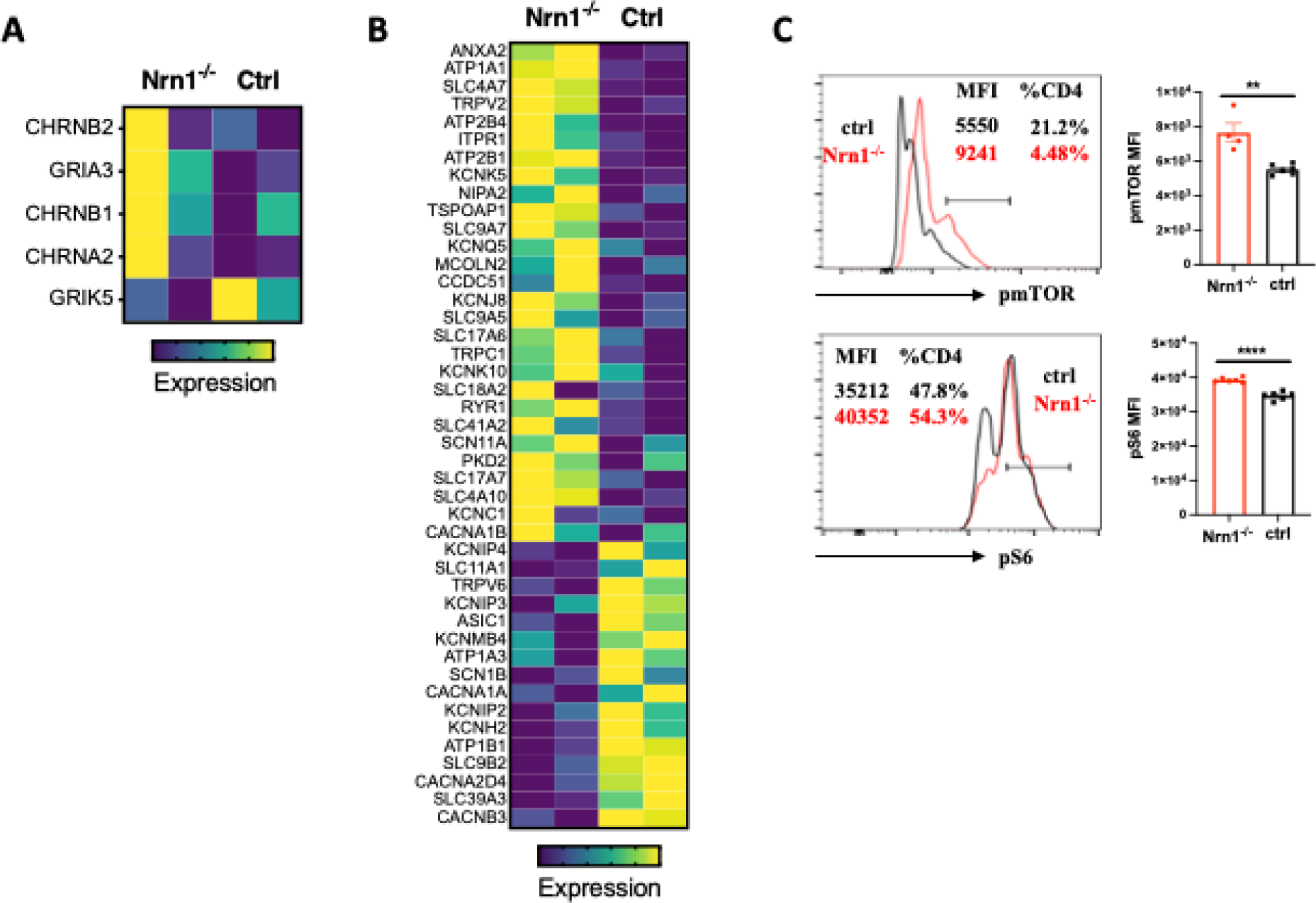
Heatmap of enriched genes in Te cells. **(A)** Differentially expressed genes in “GOMF_Neurotransmitter receptor activity involved in the regulation of postsynaptic membrane potential” gene set in Nrn1^-/-^ and ctrl Te cells. **(B)** Heatmap of differentially expressed genes in “MF_metal ion transmembrane transporter activity” in Nrn1^-/-^ and ctrl Te cells. **(C).** Detection of pmTOR and pS6 in Nrn1^-/-^ and ctrl iTreg cells. Data represents three independent experiments. **p<0.01, ****p<0.0001. Unpaired Student’s t-tests were performed.

**Figure 3-figure supplement Table 1.**
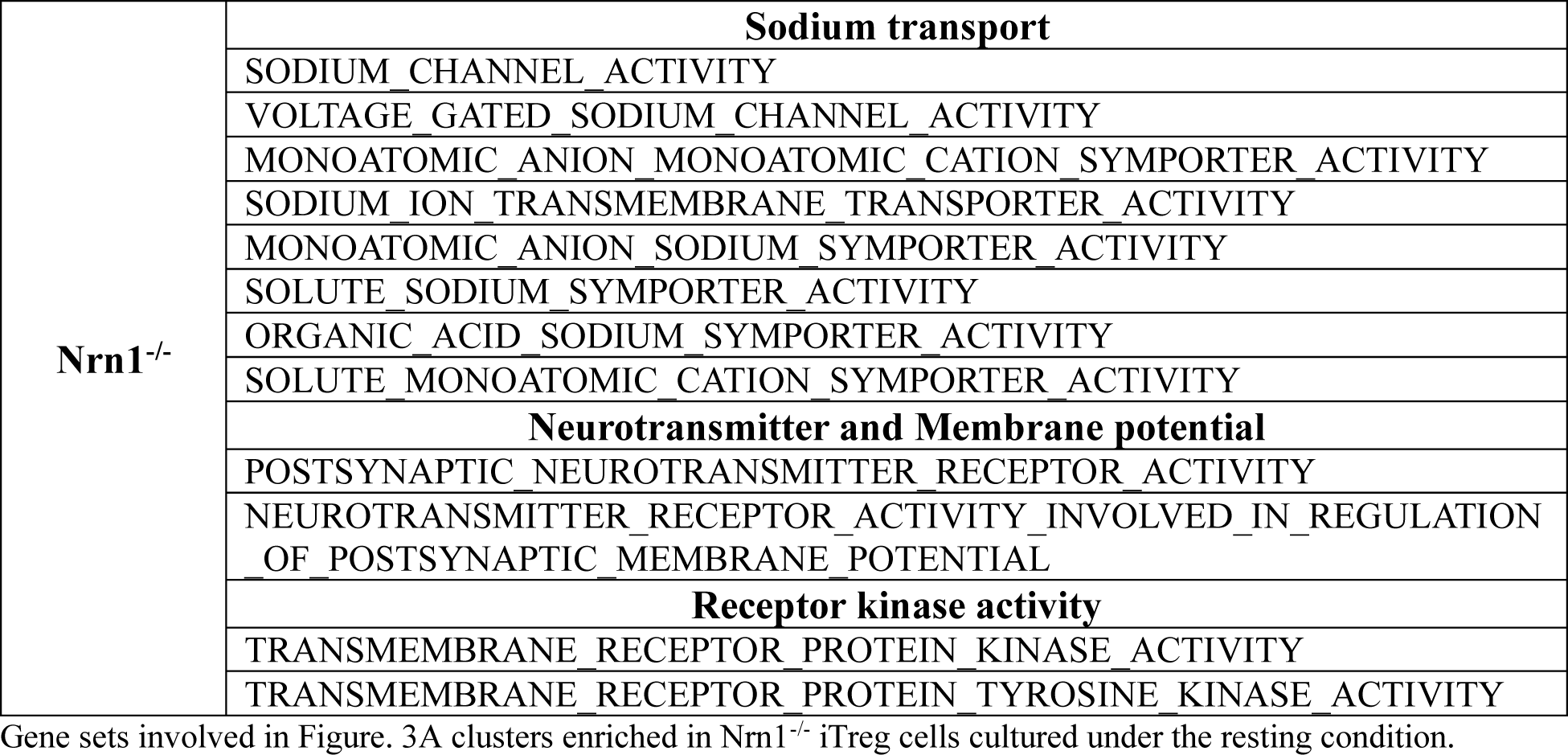
Gene sets enriched in Nrn1^-/-^ iTreg cells cultured under the resting condition.

**Figure 3-figure supplement Table 2:**
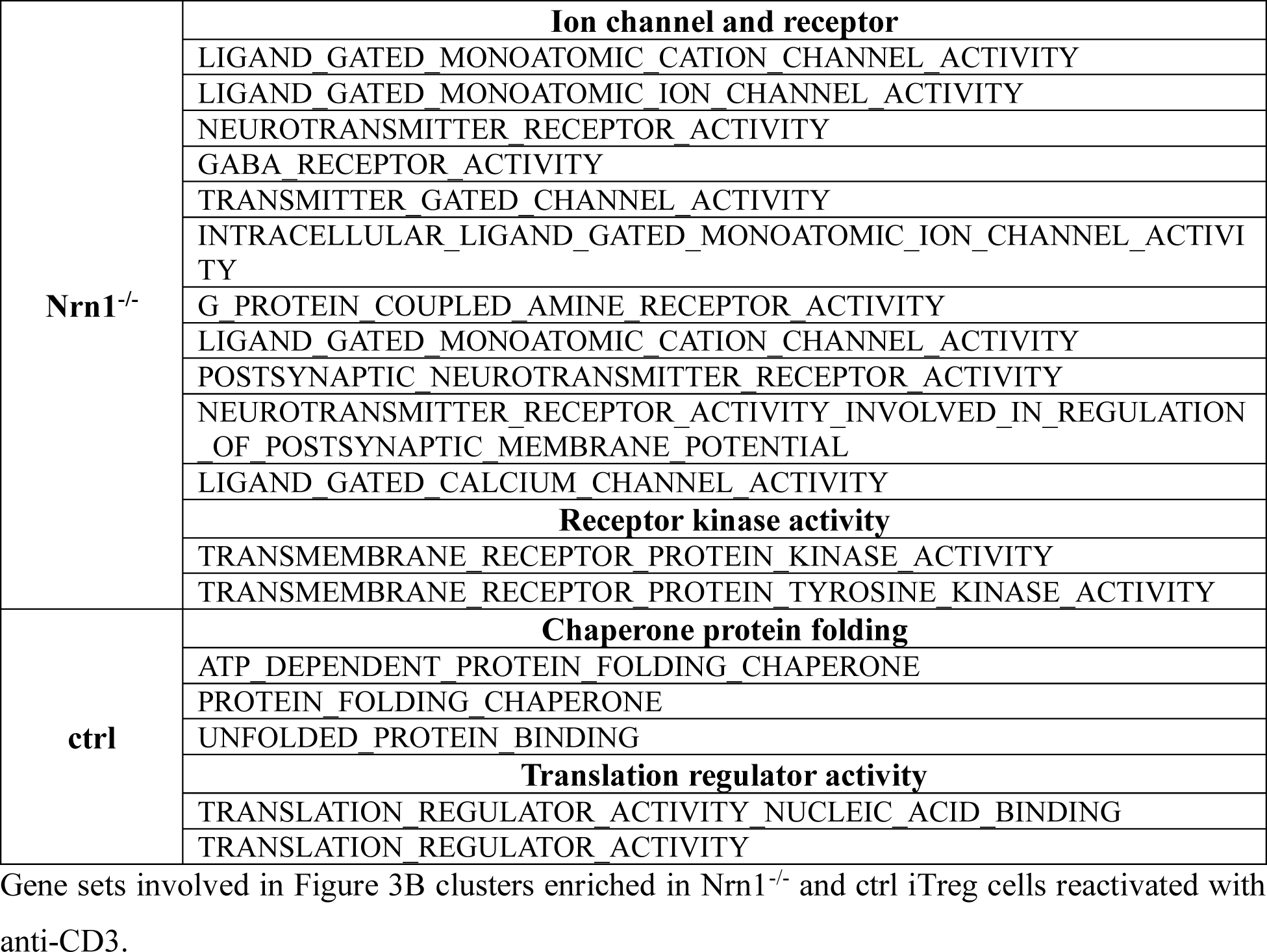
Gene sets enriched in Nrn1^-/-^ iTreg cells cultured under the reactivating condition.

**Figure 3-figure supplement Table 3:**
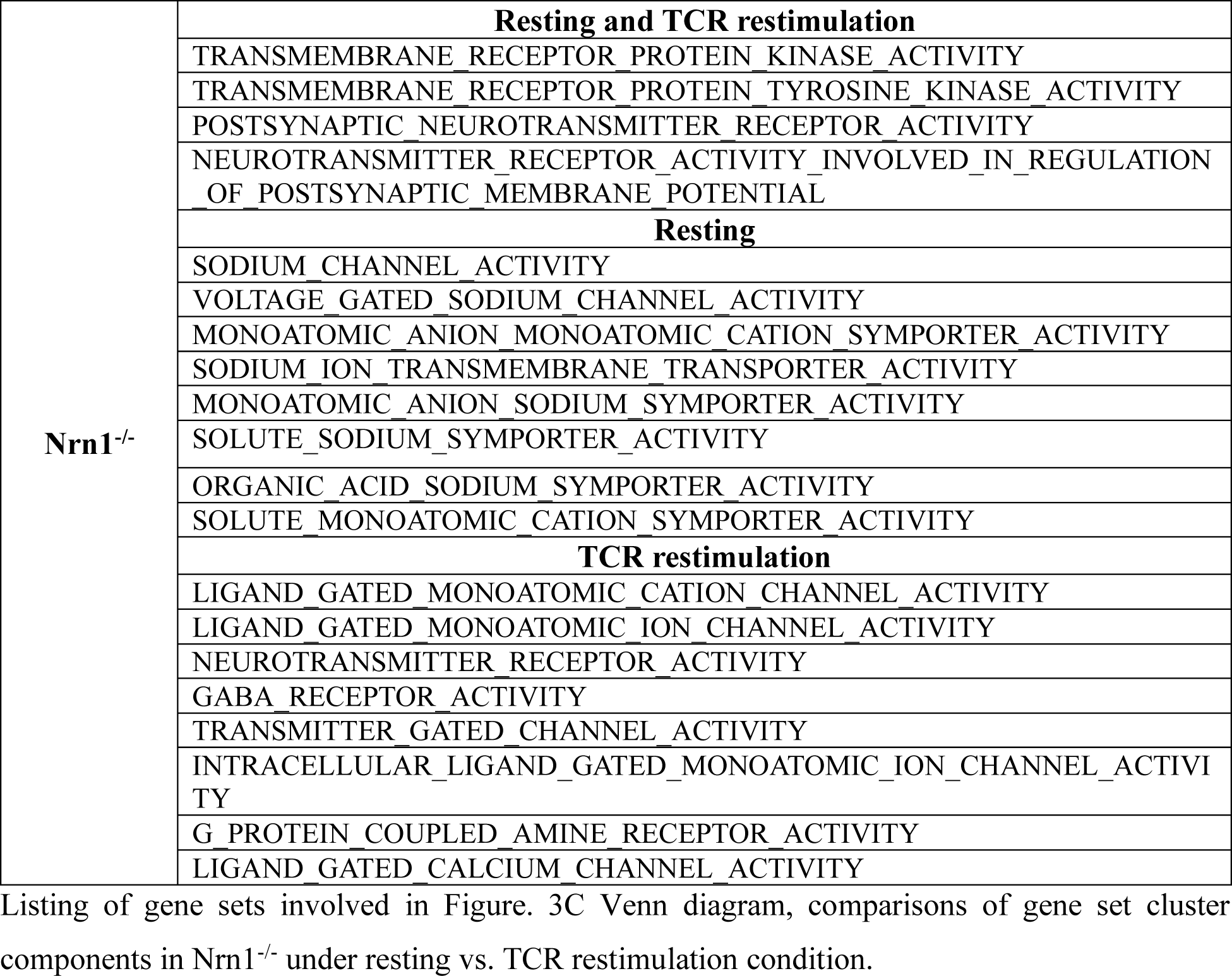
Comparison of gene sets enriched in Nrn1^-/-^ iTreg cells cultured under the resting and TCR restimulation condition.

**Figure 4-figure supplement Table 1:**
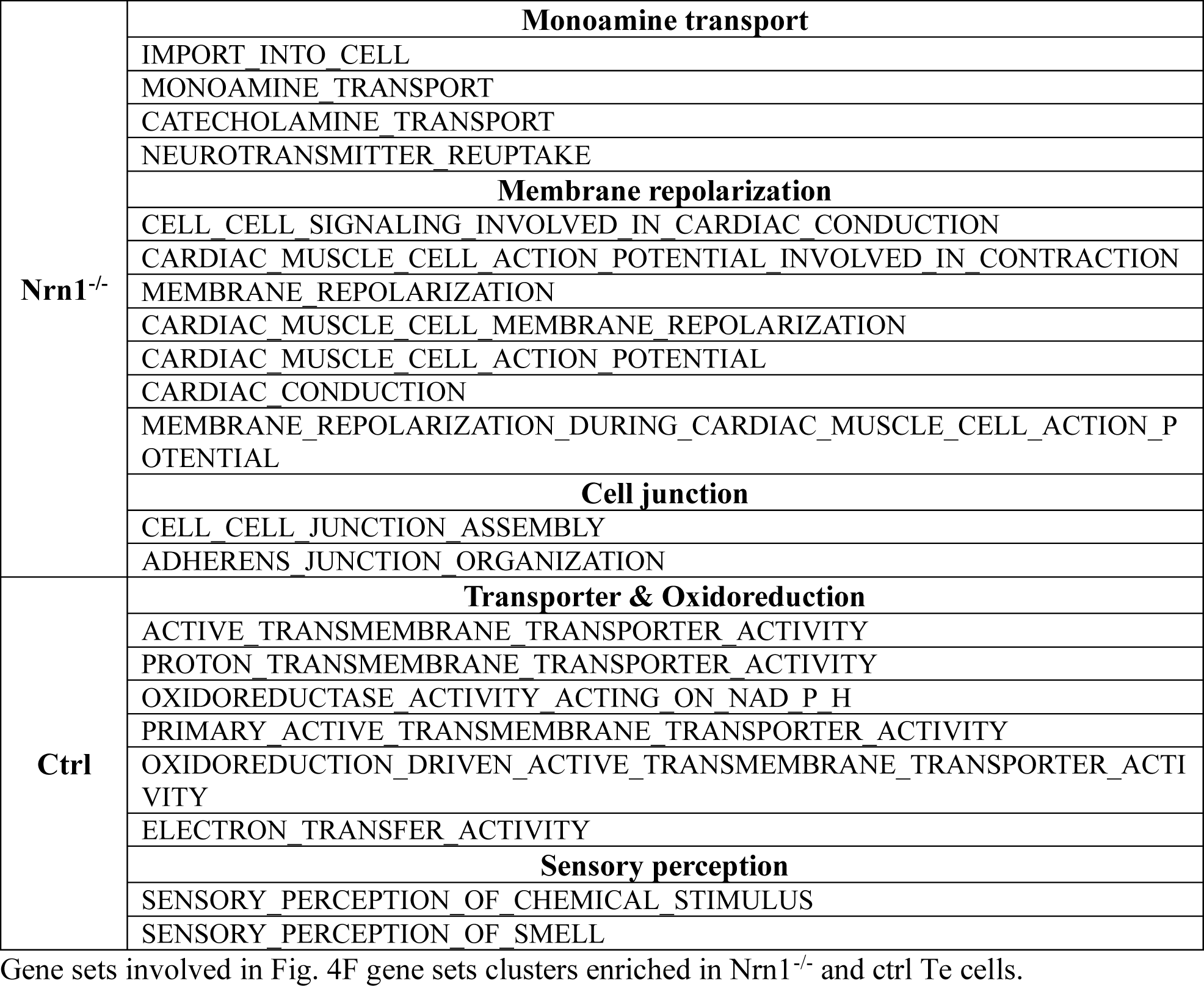
Gene sets enriched in Nrn1^-/-^ and ctrl Te cells.

## Notes

### Competing Interest Statement

CD is a co-inventor on patents licensed from JHU to BMS and Janssen and is currently an employee of Janssen Research. DMP is a consultant for Compugen, Shattuck Labs, WindMIL, Tempest, Immunai, Bristol-Myers Squibb, Amgen, Janssen, Astellas, Rockspring Capital, Immunomic, and Dracen, owns founders equity in ManaT Bio Inc., WindMIL, Trex, Jounce, Enara, Tizona, Tieza, and RAPT; and receives research funding from Compugen, Bristol-Myers Squibb, and Enara. None of the other authors have conflicting financial interests.

### Summary of Updates

We added additional experiments to confirm the following issues: 1. Amino acid (AAs) induced membrane potential change, confirming differential AA transporter expression and its potential impact on cellular MP. 2. Examine naive T cell electrical and metabolic state in the presence or absence of Nrn1 expression. 3. Examine Nrn1 blockade effect on iTreg differention. 4. Minor corrections on figure labelling, etc.

## References

Abdul Kadir, L., M. Stacey, and R. Barrett-Jolley. 2018. Emerging Roles of the Membrane Potential: Action Beyond the Action Potential. Front Physiol 9:1661.

Adler, A.J., D.W. Marsh, G.S. Yochum, J.L. Guzzo, A. Nigam, W.G. Nelson, and D.M. Pardoll. 1998. CD4+ T cell tolerance to parenchymal self-antigens requires presentation by bone marrow-derived antigen-presenting cells. The Journal of experimental medicine 187:1555–1564.

Anders, S., P.T. Pyl, and W. Huber. 2015. HTSeq--a Python framework to work with high-throughput sequencing data. Bioinformatics 31:166–169.

Babst, M. 2020. Regulation of nutrient transporters by metabolic and environmental stresses. Curr Opin Cell Biol 65:35–41.

Blackiston, D.J., K.A. McLaughlin, and M. Levin. 2009. Bioelectric controls of cell proliferation: ion channels, membrane voltage and the cell cycle. Cell Cycle 8:3527–3536.

Bohmwald, K., N.M.S. Gálvez, C.A. Andrade, V.P. Mora, J.T. Muñoz, P.A. González, C.A. Riedel, and A.M. Kalergis. 2021. Modulation of Adaptive Immunity and Viral Infections by Ion Channels. Front Physiol 12:736681.

Bonnet, C.S., S.J. Gilbert, E.J. Blain, A.S. Williams, and D.J. Mason. 2020. AMPA/kainate glutamate receptor antagonists prevent posttraumatic osteoarthritis. JCI insight 5:

Bonnet, C.S., A.S. Williams, S.J. Gilbert, A.K. Harvey, B.A. Evans, and D.J. Mason. 2015. AMPA/kainate glutamate receptors contribute to inflammation, degeneration and pain related behaviour in inflammatory stages of arthritis. Annals of the rheumatic diseases 74:242–251.

Buck, M.D., R.T. Sowell, S.M. Kaech, and E.L. Pearce. 2017. Metabolic Instruction of Immunity. Cell 169:570–586.

Cantallops, I., K. Haas, and H.T. Cline. 2000. Postsynaptic CPG15 promotes synaptic maturation and presynaptic axon arbor elaboration in vivo. Nature neuroscience 3:1004–1011.

Chapman, N.M., and H. Chi. 2022. Metabolic adaptation of lymphocytes in immunity and disease. Immunity 55:14–30.

Chappert, P., and R.H. Schwartz. 2010. Induction of T cell anergy: integration of environmental cues and infectious tolerance. Current opinion in immunology 22:552–559.

Chen, T.C., S.P. Cobbold, P.J. Fairchild, and H. Waldmann. 2004. Generation of anergic and regulatory T cells following prolonged exposure to a harmless antigen. *Journal of immunology (Baltimore*, Md*. :* 1950) 172:5900-5907.

Choi, S., and R.H. Schwartz. 2007. Molecular mechanisms for adaptive tolerance and other T cell anergy models. Seminars in immunology 19:140–152.

Cuenca, A., F. Cheng, H. Wang, J. Brayer, P. Horna, L. Gu, H. Bien, I.M. Borrello, H.I. Levitsky, and E.M. Sotomayor. 2003. Extra-lymphatic solid tumor growth is not immunologically ignored and results in early induction of antigen-specific T-cell anergy: dominant role of cross-tolerance to tumor antigens. Cancer research 63:9007–9015.

DL, K.L.a.M. 2017. Relationship between CD4 Regulatory T Cells and Anergy In Vivo. *Journal of immunology (Baltimore*, Md*. :* 1950) 198:2527.

Dobin, A., C.A. Davis, F. Schlesinger, J. Drenkow, C. Zaleski, S. Jha, P. Batut, M. Chaisson, and T.R. Gingeras. 2013. STAR: ultrafast universal RNA-seq aligner. Bioinformatics 29:15–21.

Dvorak, V., T. Wiedmer, A. Ingles-Prieto, P. Altermatt, H. Batoulis, F. Bärenz, E. Bender, D. Digles, F. Dürrenberger, L.H. Heitman, I.J. AP, D.B. Kell, S. Kickinger, D. Körzö, P. Leippe, T. Licher, V. Manolova, R. Rizzetto, F. Sassone, L. Scarabottolo, A. Schlessinger, V. Schneider, H.J. Sijben, A.L. Steck, H. Sundström, S. Tremolada, M. Wilhelm, M. Wright Muelas, D. Zindel, C.M. Steppan, and G. Superti-Furga. 2021. An Overview of Cell-Based Assay Platforms for the Solute Carrier Family of Transporters. Frontiers in pharmacology 12:722889.

ElTanbouly, M.A., and R.J. Noelle. 2021. Rethinking peripheral T cell tolerance: checkpoints across a T cell’s journey. Nat Rev Immunol 21:257–267.

Emmons-Bell, M., and I.K. Hariharan. 2021. Membrane potential regulates Hedgehog signalling in the Drosophila wing imaginal disc. EMBO reports 22:e51861.

Feng, Y., A. Arvey, T. Chinen, J. van der Veeken, G. Gasteiger, and A.Y. Rudensky. 2014. Control of the inheritance of regulatory T cell identity by a cis element in the Foxp3 locus. Cell 158:749–763.

Floess, S., J. Freyer, C. Siewert, U. Baron, S. Olek, J. Polansky, K. Schlawe, H.D. Chang, T. Bopp, E. Schmitt, S. Klein-Hessling, E. Serfling, A. Hamann, and J. Huehn. 2007. Epigenetic control of the foxp3 locus in regulatory T cells. PLoS biology 5:e38.

Fujino, T., J.H. Leslie, R. Eavri, J.L. Chen, W.C. Lin, G.H. Flanders, E. Borok, T.L. Horvath, and E. Nedivi. 2011. CPG15 regulates synapse stability in the developing and adult brain. Genes & development 25:2674–2685.

Geltink, R.I.K., R.L. Kyle, and E.L. Pearce. 2018. Unraveling the Complex Interplay Between T Cell Metabolism and Function. Annual review of immunology 36:461–488.

Gonzalez-Figueroa, P., J.A. Roco, I. Papa, L. Núñez Villacís, M. Stanley, M.A. Linterman, A. Dent, P.F. Canete, and C.G. Vinuesa. 2021. Follicular regulatory T cells produce neuritin to regulate B cells. Cell

Hamill, M.J., R. Afeyan, M.V. Chakravarthy, and T. Tramontin. 2020. Endogenous Metabolic Modulators: Emerging Therapeutic Potential of Amino Acids. iScience 23:101628.

Huang, C.T., D.L. Huso, Z. Lu, T. Wang, G. Zhou, E.P. Kennedy, C.G. Drake, D.J. Morgan, L.A. Sherman, A.D. Higgins, D.M. Pardoll, and A.J. Adler. 2003. CD4+ T cells pass through an effector phase during the process of in vivo tolerance induction. Journal of immunology (Baltimore, Md. : 1950) 170:3945-3953.

Huang, C.T., C.J. Workman, D. Flies, X. Pan, A.L. Marson, G. Zhou, E.L. Hipkiss, S. Ravi, J. Kowalski, H.I. Levitsky, J.D. Powell, D.M. Pardoll, C.G. Drake, and D.A. Vignali. 2004. Role of LAG-3 in regulatory T cells. Immunity 21:503–513.

Javaherian, A., and H.T. Cline. 2005. Coordinated motor neuron axon growth and neuromuscular synaptogenesis are promoted by CPG15 in vivo. Neuron 45:505–512.

Joesch, C., E. Guevarra, S.P. Parel, A. Bergner, P. Zbinden, D. Konrad, and H. Albrecht. 2008. Use of FLIPR membrane potential dyes for validation of high-throughput screening with the FLIPR and microARCS technologies: identification of ion channel modulators acting on the GABA(A) receptor. J Biomol Screen 13:218–228.

Kalekar, L.A., S.E. Schmiel, S.L. Nandiwada, W.Y. Lam, L.O. Barsness, N. Zhang, G.L. Stritesky, D. Malhotra, K.E. Pauken, J.L. Linehan, M.G. O’Sullivan, B.T. Fife, K.A. Hogquist, M.K. Jenkins, and D.L. Mueller. 2016. CD4(+) T cell anergy prevents autoimmunity and generates regulatory T cell precursors. Nature immunology 17:304–314.

Kiefer, H., A.J. Blume, and H.R. Kaback. 1980. Membrane potential changes during mitogenic stimulation of mouse spleen lymphocytes. Proceedings of the National Academy of Sciences of the United States of America 77:2200–2204.

Kim, J.M., J.P. Rasmussen, and A.Y. Rudensky. 2007. Regulatory T cells prevent catastrophic autoimmunity throughout the lifespan of mice. Nature immunology 8:191–197.

Kuczma, M.P., E.A. Szurek, A. Cebula, V.L. Ngo, M. Pietrzak, P. Kraj, T.L. Denning, and L. Ignatowicz. 2021. Self and microbiota-derived epitopes induce CD4(+) T cell anergy and conversion into CD4(+)Foxp3(+) regulatory cells. Mucosal Immunol 14:443–454.

Levin, M. 2021. Bioelectric signaling: Reprogrammable circuits underlying embryogenesis, regeneration, and cancer. Cell 184:1971–1989.

Li, X., Y. Liang, M. LeBlanc, C. Benner, and Y. Zheng. 2014. Function of a Foxp3 cis-element in protecting regulatory T cell identity. Cell 158:734–748.

Lim, D.G., Y.H. Park, S.E. Kim, S.H. Jeong, and S.C. Kim. 2013. Diagnostic value of tolerance- related gene expression measured in the recipient alloantigen-reactive T cell fraction. *Clinical immunology (Orlando*, Fla.) 148:219–226.

Liu, G.Y., and D.M. Sabatini. 2020. mTOR at the nexus of nutrition, growth, ageing and disease. Nat Rev Mol Cell Biol 21:183–203.

Long, L., J. Wei, S.A. Lim, J.L. Raynor, H. Shi, J.P. Connelly, H. Wang, C. Guy, B. Xie, N.M. Chapman, G. Fu, Y. Wang, H. Huang, W. Su, J. Saravia, I. Risch, Y.D. Wang, Y. Li, M. Niu, Y. Dhungana, A. Kc, P. Zhou, P. Vogel, J. Yu, S.M. Pruett-Miller, J. Peng, and H. Chi. 2021. CRISPR screens unveil signal hubs for nutrient licensing of T cell immunity. Nature 600:308–313.

Love, M.I., W. Huber, and S. Anders. 2014. Moderated estimation of fold change and dispersion for RNA-seq data with DESeq2. Genome biology 15:550.

Ma, Y., K. Poole, J. Goyette, and K. Gaus. 2017. Introducing Membrane Charge and Membrane Potential to T Cell Signaling. Frontiers in immunology 8:1513.

Martinez, R.J., N. Zhang, S.R. Thomas, S.L. Nandiwada, M.K. Jenkins, B.A. Binstadt, and D.L. Mueller. 2012. Arthritogenic self-reactive CD4+ T cells acquire an FR4hiCD73hi anergic state in the presence of Foxp3+ regulatory T cells. *Journal of immunology (Baltimore*, Md*. :* 1950) 188:170-181.

McNearney, T., B.A. Baethge, S. Cao, R. Alam, J.R. Lisse, and K.N. Westlund. 2004. Excitatory amino acids, TNF-alpha, and chemokine levels in synovial fluids of patients with active arthropathies. Clinical and experimental immunology 137:621–627.

Mercadante, E.R., and U.M. Lorenz. 2016. Breaking Free of Control: How Conventional T Cells Overcome Regulatory T Cell Suppression. Frontiers in immunology 7:193.

Miller, S.D., W.J. Karpus, and T.S. Davidson. 2007. Experimental Autoimmune Encephalomyelitis in the Mouse. Current protocols in immunology CHAPTER:Unit-15 11.

Monroe, J.G., and J.C. Cambier. 1983. B cell activation. I. Anti-immunoglobulin-induced receptor cross-linking results in a decrease in the plasma membrane potential of murine B lymphocytes. The Journal of experimental medicine 157:2073–2086.

Nedivi, E., G.Y. Wu, and H.T. Cline. 1998. Promotion of dendritic growth by CPG15, an activity- induced signaling molecule. *Science (New York*, N.Y.) 281:1863–1866.

Nik, A.M., B. Pressly, V. Singh, S. Antrobus, S. Hulsizer, M.A. Rogawski, H. Wulff, and I.N. Pessah. 2017. Rapid Throughput Analysis of GABA(A) Receptor Subtype Modulators and Blockers Using DiSBAC(1)(3) Membrane Potential Red Dye. Mol Pharmacol 92:88–99.

Nystrom, S.N., D. Bourges, S. Garry, E.M. Ross, I.R. van Driel, and P.A. Gleeson. 2014. Transient Treg-cell depletion in adult mice results in persistent self-reactive CD4(+) T-cell responses. European journal of immunology 44:3621–3631.

Olenchock, B.A., J.C. Rathmell, and M.G. Vander Heiden. 2017. Biochemical Underpinnings of Immune Cell Metabolic Phenotypes. Immunity 46:703–713.

Opejin, A., A. Surnov, Z. Misulovin, M. Pherson, C. Gross, C.A. Iberg, I. Fallahee, J. Bourque, D. Dorsett, and D. Hawiger. 2020. A Two-Step Process of Effector Programming Governs CD4(+) T Cell Fate Determination Induced by Antigenic Activation in the Steady State. Cell reports 33:108424.

Pandya, N.J., C. Seeger, N. Babai, M.A. Gonzalez-Lozano, V. Mack, J.C. Lodder, Y. Gouwenberg, H.D. Mansvelder, U.H. Danielson, K.W. Li, M. Heine, S. Spijker, R. Frischknecht, and A.B. Smit. 2018. Noelin1 Affects Lateral Mobility of Synaptic AMPA Receptors. Cell reports 24:1218–1230.

Peng, M., and M.O. Li. 2023. Metabolism along the life journey of T cells. Life Metab 2:

Plitas, G., C. Konopacki, K. Wu, P.D. Bos, M. Morrow, E.V. Putintseva, D.M. Chudakov, and A.Y. Rudensky. 2016. Regulatory T Cells Exhibit Distinct Features in Human Breast Cancer. Immunity 45:1122–1134.

Putz, U., C. Harwell, and E. Nedivi. 2005. Soluble CPG15 expressed during early development rescues cortical progenitors from apoptosis. Nature neuroscience 8:322–331.

Ramirez, G.A., L.A. Coletto, C. Sciorati, E.P. Bozzolo, P. Manunta, P. Rovere-Querini, and A.A. Manfredi. 2018. Ion Channels and Transporters in Inflammation: Special Focus on TRP Channels and TRPC6. Cells 7:

Safford, M., S. Collins, M.A. Lutz, A. Allen, C.T. Huang, J. Kowalski, A. Blackford, M.R. Horton, C. Drake, R.H. Schwartz, and J.D. Powell. 2005. Egr-2 and Egr-3 are negative regulators of T cell activation. Nature immunology 6:472–480.

Salmond, R.J. 2018. mTOR Regulation of Glycolytic Metabolism in T Cells. Frontiers in cell and developmental biology 6:122.

Saravia, J., J.L. Raynor, N.M. Chapman, S.A. Lim, and H. Chi. 2020. Signaling networks in immunometabolism. Cell research 30:328–342.

Sarchielli, P., M. Di Filippo, A. Candeliere, D. Chiasserini, A. Mattioni, S. Tenaglia, M. Bonucci, and P. Calabresi. 2007. Expression of ionotropic glutamate receptor GLUR3 and effects of glutamate on MBP- and MOG-specific lymphocyte activation and chemotactic migration in multiple sclerosis patients. Journal of neuroimmunology 188:146–158.

Schietinger, A., J.J. Delrow, R.S. Basom, J.N. Blattman, and P.D. Greenberg. 2012. Rescued tolerant CD8 T cells are preprogrammed to reestablish the tolerant state. *Science (New York*, N.Y.) 335:723–727.

Schietinger, A., M. Philip, V.E. Krisnawan, E.Y. Chiu, J.J. Delrow, R.S. Basom, P. Lauer, D.G. Brockstedt, S.E. Knoblaugh, G.J. Hammerling, T.D. Schell, N. Garbi, and P.D. Greenberg. 2016. Tumor-Specific T Cell Dysfunction Is a Dynamic Antigen-Driven Differentiation Program Initiated Early during Tumorigenesis. Immunity 45:389–401.

Schwenk, J., N. Harmel, A. Brechet, G. Zolles, H. Berkefeld, C.S. Muller, W. Bildl, D. Baehrens, B. Huber, A. Kulik, N. Klocker, U. Schulte, and B. Fakler. 2012. High-resolution proteomics unravel architecture and molecular diversity of native AMPA receptor complexes. Neuron 74:621–633.

Shannon, P., A. Markiel, O. Ozier, N.S. Baliga, J.T. Wang, D. Ramage, N. Amin, B. Schwikowski, and T. Ideker. 2003. Cytoscape: a software environment for integrated models of biomolecular interaction networks. Genome Res 13:2498–2504.

Shimada, T., T. Yoshida, and K. Yamagata. 2016. Neuritin Mediates Activity-Dependent Axonal Branch Formation in Part via FGF Signaling. The Journal of neuroscience : the official journal of the Society for Neuroscience 36:4534–4548.

Shin, D.S., A. Jordan, S. Basu, R.M. Thomas, S. Bandyopadhyay, E.F. de Zoeten, A.D. Wells, and F. Macian. 2014. Regulatory T cells suppress CD4+ T cells through NFAT-dependent transcriptional mechanisms. EMBO reports 15:991–999.

Silva Morales, M., and D. Mueller. 2018. Anergy into T regulatory cells: an integration of metabolic cues and epigenetic changes at the Foxp3 conserved non-coding sequence 2. F1000Research 7:

Sinclair, L.V., J. Rolf, E. Emslie, Y.B. Shi, P.M. Taylor, and D.A. Cantrell. 2013. Control of amino-acid transport by antigen receptors coordinates the metabolic reprogramming essential for T cell differentiation. Nature immunology 14:500–508.

Singer, M., C. Wang, L. Cong, N.D. Marjanovic, M.S. Kowalczyk, H. Zhang, J. Nyman, K. Sakuishi, S. Kurtulus, D. Gennert, J. Xia, J.Y. Kwon, J. Nevin, R.H. Herbst, I. Yanai, O. Rozenblatt-Rosen, V.K. Kuchroo, A. Regev, and A.C. Anderson. 2016. A Distinct Gene Module for Dysfunction Uncoupled from Activation in Tumor-Infiltrating T Cells. Cell 166:1500–1511.e1509.

Subramanian, A., P. Tamayo, V.K. Mootha, S. Mukherjee, B.L. Ebert, M.A. Gillette, A. Paulovich, S.L. Pomeroy, T.R. Golub, E.S. Lander, and J.P. Mesirov. 2005. Gene set enrichment analysis: a knowledge-based approach for interpreting genome-wide expression profiles. Proceedings of the National Academy of Sciences of the United States of America 102:15545–15550.

Subramanian, J., K. Michel, M. Benoit, and E. Nedivi. 2019. CPG15/Neuritin Mimics Experience in Selecting Excitatory Synapses for Stabilization by Facilitating PSD95 Recruitment. Cell reports 28:1584–1595.e1585.

Sullivan, M.R., L.V. Danai, C.A. Lewis, S.H. Chan, D.Y. Gui, T. Kunchok, E.A. Dennstedt, M.G. Vander Heiden, and A. Muir. 2019. Quantification of microenvironmental metabolites in murine cancers reveals determinants of tumor nutrient availability. eLife 8:

Sundelacruz, S., M. Levin, and D.L. Kaplan. 2009. Role of membrane potential in the regulation of cell proliferation and differentiation. Stem Cell Rev Rep 5:231–246.

Vahl, J.C., C. Drees, K. Heger, S. Heink, J.C. Fischer, J. Nedjic, N. Ohkura, H. Morikawa, H. Poeck, S. Schallenberg, D. Riess, M.Y. Hein, T. Buch, B. Polic, A. Schonle, R. Zeiser, A. Schmitt-Graff, K. Kretschmer, L. Klein, T. Korn, S. Sakaguchi, and M. Schmidt-Supprian. 2014. Continuous T cell receptor signals maintain a functional regulatory T cell pool. Immunity 41:722–736.

Vanasek, T.L., S.L. Nandiwada, M.K. Jenkins, and D.L. Mueller. 2006. CD25+Foxp3+ regulatory T cells facilitate CD4+ T cell clonal anergy induction during the recovery from lymphopenia. *Journal of immunology (Baltimore*, Md*. :* 1950) 176:5880-5889.

Wang, Y., A. Tao, M. Vaeth, and S. Feske. 2020. Calcium regulation of T cell metabolism. Current opinion in physiology 17:207–223.

Whiteaker, K.L., S.M. Gopalakrishnan, D. Groebe, C.C. Shieh, U. Warrior, D.J. Burns, M.J. Coghlan, V.E. Scott, and M. Gopalakrishnan. 2001. Validation of FLIPR membrane potential dye for high throughput screening of potassium channel modulators. J Biomol Screen 6:305–312.

Workman, C.J., L.W. Collison, M. Bettini, M.R. Pillai, J.E. Rehg, and D.A. Vignali. 2011. In vivo Treg suppression assays. *Methods in molecular biology (Clifton*, N.J.) 707:119–156.

Yao, J.J., X.F. Gao, C.W. Chow, X.Q. Zhan, C.L. Hu, and Y.A. Mei. 2012. Neuritin activates insulin receptor pathway to up-regulate Kv4.2-mediated transient outward K+ current in rat cerebellar granule neurons. The Journal of biological chemistry 287:41534–41545.

Yu, W., Z. Wang, X. Yu, Y. Zhao, Z. Xie, K. Zhang, Z. Chi, S. Chen, T. Xu, D. Jiang, X. Guo, M. Li, J. Zhang, H. Fang, D. Yang, Y. Guo, X. Yang, X. Zhang, Y. Wu, W. Yang, and D. Wang. 2022. Kir2.1-mediated membrane potential promotes nutrient acquisition and inflammation through regulation of nutrient transporters. Nature communications 13:3544.

Zha, Y., R. Marks, A.W. Ho, A.C. Peterson, S. Janardhan, I. Brown, K. Praveen, S. Stang, J.C. Stone, and T.F. Gajewski. 2006. T cell anergy is reversed by active Ras and is regulated by diacylglycerol kinase-alpha. Nature immunology 7:1166–1173.

Zheng, Y., G.M. Delgoffe, C.F. Meyer, W. Chan, and J.D. Powell. 2009. Anergic T cells are metabolically anergic. *Journal of immunology (Baltimore*, Md*. :* 1950) 183:6095-6101.

Zheng, Y., S. Josefowicz, A. Chaudhry, X.P. Peng, K. Forbush, and A.Y. Rudensky. 2010. Role of conserved non-coding DNA elements in the Foxp3 gene in regulatory T-cell fate. Nature 463:808–812.

Zhou, S., and J. Zhou. 2014. Neuritin, a neurotrophic factor in nervous system physiology. Current medicinal chemistry 21:1212–1219.

Zhou, Y., C.O. Wong, K.J. Cho, D. van der Hoeven, H. Liang, D.P. Thakur, J. Luo, M. Babic, K.E. Zinsmaier, M.X. Zhu, H. Hu, K. Venkatachalam, and J.F. Hancock. 2015. SIGNAL TRANSDUCTION. Membrane potential modulates plasma membrane phospholipid dynamics and K-Ras signaling. *Science (New York*, N.Y.) 349:873–876.

Zito, A., D. Cartelli, G. Cappelletti, A. Cariboni, W. Andrews, J. Parnavelas, A. Poletti, and M. Galbiati. 2014. Neuritin 1 promotes neuronal migration. Brain Structure and Function 219:105–118.

