## supplemental figures for "Neurotrophic factor Neuritin modulates T cell electrical and metabolic state for the balance of tolerance and immunity"

**Figure supplements**

**Hong Yu *et al*.**

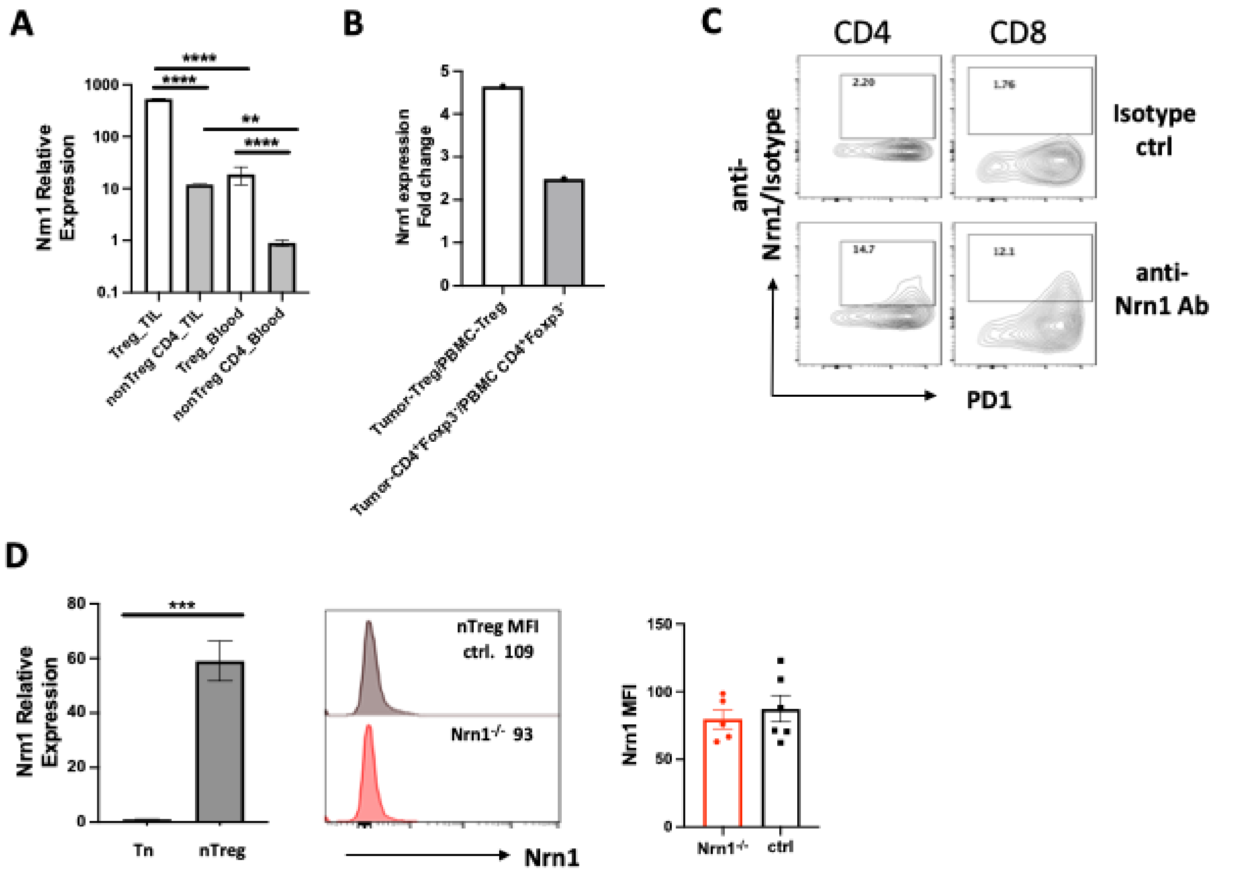


**Figure 1-figure supplement 1. Nrn1 expression in T cells from tumor environment and during early T cell activation. (A-B).** Nrn1 expression in tumor infiltrates. (A) Comparison of Nrn1 expression by qRT-PCR among Tregs and non-Treg $CD44^{hi}CD4^{+}$ cells recovered either from B16 melanoma infiltrates or from peripheral blood of Foxp3DTRgfp mice bearing subcutaneous B16 melanomas. **(B)** Comparison of Nrn1 expression in breast tumor-infiltrating Treg (T-Treg) and Te (T-$CD4^{+}Foxp3^{-}$) cells vs. peripheral blood Treg (P-Treg) and Te (PBMC $CD4^{+}Foxp3^{-}$) cells. Data derived from the “Regulatory T Cells Exhibit Distinct Features in Human Breast Cancer” report (Plitas et al., 2016). **(C)** Nrn1 cell surface detection on day 2 activated CD4 and CD8 cells by flow cytometry. **(D).** Detection of Nrn1 expression in nTreg cells by qRT-PCR and cell surface Nrn1 staining.

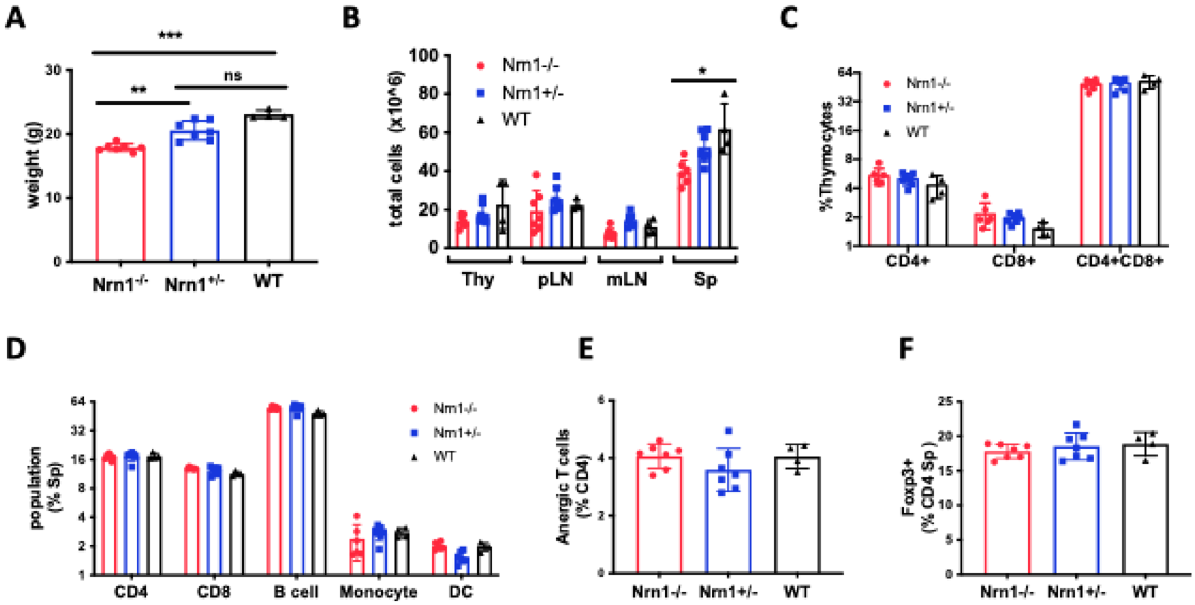


**Figure 1-figure supplement 2. Nrn1$^{-/-}$ mice body weight and immune cell profile analysis compared to Nrn1$^{+/-}$ and WT mice. (A)** Average body weight of 10-12 wk old age and sex-matched Nrn1$^{-/-}$, Nrn1$^{+/-}$ and WT mice. **(B)** Thymus and peripheral lymphoid tissue total cell count. **(C-D)** Immune cell frequencies in the thymus and spleen. **(E)** Proportion of CD4$^{+}$Foxp3$^{-}$ CD44$^{+}$FR4$^{hi}$ CD73$^{hi}$ anergic T cells among splenocytes CD4 cell population. **(F)** FOXP3$^{+}$cell frequency among CD4 cells in the spleen. The immune profile assessment used n>3 mice/group of Nrn1$^{-/-}$ and control mice. *P* values were calculated by one-way analysis of variance (ANOVA). **P*<0.05. ***P*<0.01.

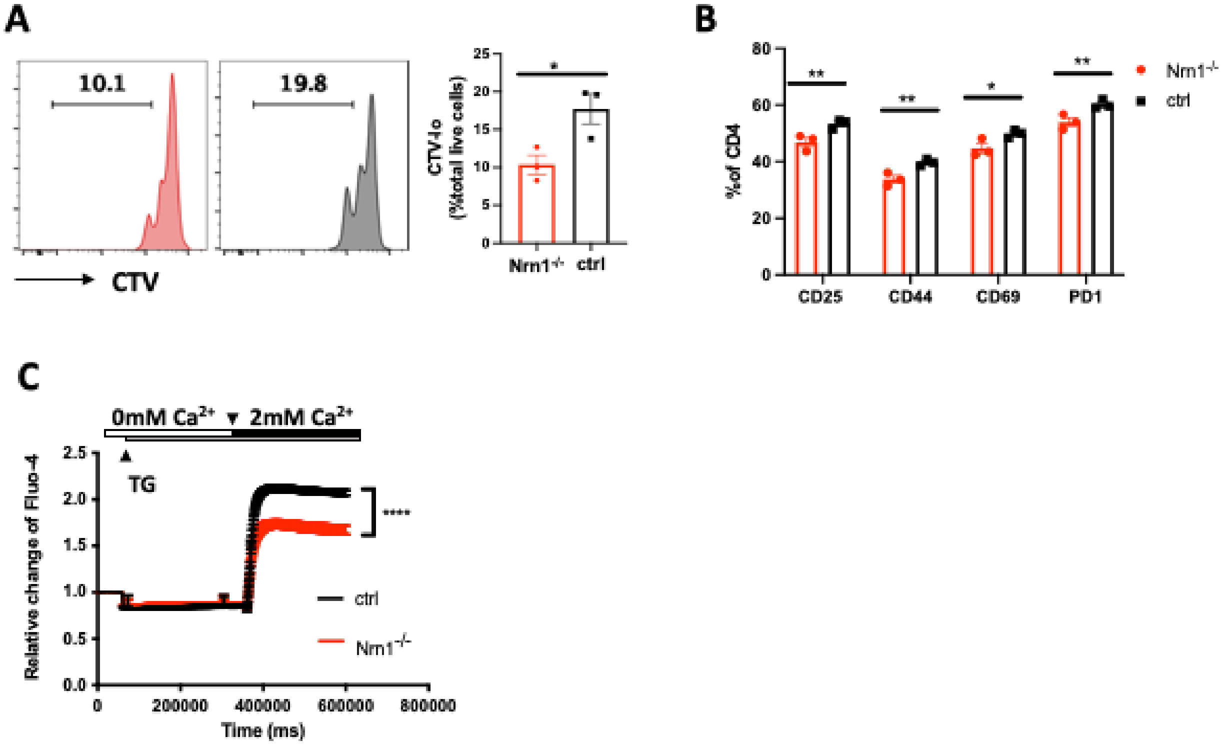


**Figure 1-figure supplement 3. Compromised T cell activation in Nrn1$^{-/-}$ cells. (A)** Cell tracker dye violet (CTV) dilution in Nrn1$^{-/-}$ or ctrl CD4 cells after stimulation with plate-bound aCD3 (5ug/ml) and soluble aCD28 (2ug/ml); **(B)** Cell surface activation markers CD25, CD44, CD69, and PD1 expression day 2 after naïve CD4$^{+}$ cell activation. Unpaired student's t-test, *p<0.05, **p<0.01. Data represent three independent experiments. **(C)** Store-operated Ca$^{++}$ entry (SOCE) was examined on day 2 activated CD4$^{+}$ cells labeled with Fluo-4 dye. Representative graph and mean $\pm$ SEM of SOCE induced by CD4 cell stimulation with 1uM thapsigargin (TG) in Ca$^{++}$ free HBSS (0mM Ca$^{++}$) followed by addition of 2mM Ca$^{++}$. Representative graphs of Ca$^{++}$ influx from three independent experiments (****p<0.0001).

**Figure 3-figure supplement Table 1. Gene sets enriched in Nrn1$^{-/-}$ iTreg cells cultured under the resting condition.**

| | |
|---|---|
| **Nrn1$^{-/-}$** | **Sodium transport** |
| | SODIUM_CHANNEL_ACTIVITY |
| | VOLTAGE_GATED_SODIUM_CHANNEL_ACTIVITY |
| | MONOATOMIC_ANION_MONOATOMIC_CATION_SYMPORTER_ACTIVITY |
| | SODIUM_ION_TRANSMEMBRANE_TRANSPORTER_ACTIVITY |
| | MONOATOMIC_ANION_SODIUM_SYMPORTER_ACTIVITY |
| | SOLUTE_SODIUM_SYMPORTER_ACTIVITY |
| | ORGANIC_ACID_SODIUM_SYMPORTER_ACTIVITY |
| | SOLUTE_MONOATOMIC_CATION_SYMPORTER_ACTIVITY |
| | **Neurotransmitter and Membrane potential** |
| | POSTSYNAPTIC_NEUROTRANSMITTER_RECEPTOR_ACTIVITY |
| | NEUROTRANSMITTER_RECEPTOR_ACTIVITY_INVOLVED_IN_REGULATION_OF_POSTSYNAPTIC_MEMBRANE_POTENTIAL |
| | **Receptor kinase activity** |
| | TRANSMEMBRANE_RECEPTOR_PROTEIN_KINASE_ACTIVITY |
| | TRANSMEMBRANE_RECEPTOR_PROTEIN_TYROSINE_KINASE_ACTIVITY |

Gene sets involved in Figure. 3A clusters enriched in Nrn1$^{-/-}$ iTreg cells cultured under the resting condition.

**Figure 3-figure supplement Table 2: Gene sets enriched in Nrn1$^{-/-}$ iTreg cells cultured under the reactivating condition.**

| | |
|---|---|
| **Nrn1$^{-/-}$** | **Ion channel and receptor** |
| | LIGAND_GATED_MONOATOMIC_CATION_CHANNEL_ACTIVITY |
| | LIGAND_GATED_MONOATOMIC_ION_CHANNEL_ACTIVITY |
| | NEUROTRANSMITTER_RECEPTOR_ACTIVITY |
| | GABA_RECEPTOR_ACTIVITY |
| | TRANSMITTER_GATED_CHANNEL_ACTIVITY |
| | INTRACELLULAR_LIGAND_GATED_MONOATOMIC_ION_CHANNEL_ACTIVITY |
| | G_PROTEIN_COUPLED_AMINE_RECEPTOR_ACTIVITY |
| | LIGAND_GATED_MONOATOMIC_CATION_CHANNEL_ACTIVITY |
| | POSTSYNAPTIC_NEUROTRANSMITTER_RECEPTOR_ACTIVITY |
| | NEUROTRANSMITTER_RECEPTOR_ACTIVITY_INVOLVED_IN_REGULATION_OF_POSTSYNAPTIC_MEMBRANE_POTENTIAL |
| | LIGAND_GATED_CALCIUM_CHANNEL_ACTIVITY |
| | **Receptor kinase activity** |
| | TRANSMEMBRANE_RECEPTOR_PROTEIN_KINASE_ACTIVITY |
| | TRANSMEMBRANE_RECEPTOR_PROTEIN_TYROSINE_KINASE_ACTIVITY |
| **ctrl** | **Chaperone protein folding** |
| | ATP_DEPENDENT_PROTEIN_FOLDING_CHAPERONE |
| | PROTEIN_FOLDING_CHAPERONE |
| | UNFOLDED_PROTEIN_BINDING |
| | **Translation regulator activity** |
| | TRANSLATION_REGULATOR_ACTIVITY_NUCLEIC_ACID_BINDING |
| | TRANSLATION_REGULATOR_ACTIVITY |

Gene sets involved in Figure 3B clusters enriched in Nrn1$^{-/-}$ and ctrl iTreg cells reactivated with anti-CD3.

**Figure 3-figure supplement Table 3: Comparison of gene sets enriched in Nrn1$^{-/-}$ iTreg cells cultured under the resting and TCR restimulation condition.**

| | |
|---|---|
| **Nrn1$^{-/-}$** | **Resting and TCR restimulation** |
| | TRANSMEMBRANE_RECEPTOR_PROTEIN_KINASE_ACTIVITY |
| | TRANSMEMBRANE_RECEPTOR_PROTEIN_TYROSINE_KINASE_ACTIVITY |
| | POSTSYNAPTIC_NEUROTRANSMITTER_RECEPTOR_ACTIVITY |
| | NEUROTRANSMITTER_RECEPTOR_ACTIVITY_INVOLVED_IN_REGULATION_OF_POSTSYNAPTIC_MEMBRANE_POTENTIAL |
| | **Resting** |
| | SODIUM_CHANNEL_ACTIVITY |
| | VOLTAGE_GATED_SODIUM_CHANNEL_ACTIVITY |
| | MONOATOMIC_ANION_MONOATOMIC_CATION_SYMPORTER_ACTIVITY |
| | SODIUM_ION_TRANSMEMBRANE_TRANSPORTER_ACTIVITY |
| | MONOATOMIC_ANION_SODIUM_SYMPORTER_ACTIVITY |
| | SOLUTE_SODIUM_SYMPORTER_ACTIVITY |
| | ORGANIC_ACID_SODIUM_SYMPORTER_ACTIVITY |
| | SOLUTE_MONOATOMIC_CATION_SYMPORTER_ACTIVITY |
| | **TCR restimulation** |
| | LIGAND_GATED_MONOATOMIC_CATION_CHANNEL_ACTIVITY |
| | LIGAND_GATED_MONOATOMIC_ION_CHANNEL_ACTIVITY |
| | NEUROTRANSMITTER_RECEPTOR_ACTIVITY |
| | GABA_RECEPTOR_ACTIVITY |
| | TRANSMITTER_GATED_CHANNEL_ACTIVITY |
| | INTRACELLULAR_LIGAND_GATED_MONOATOMIC_ION_CHANNEL_ACTIVITY |
| | G_PROTEIN_COUPLED_AMINE_RECEPTOR_ACTIVITY |
| | LIGAND_GATED_CALCIUM_CHANNEL_ACTIVITY |

Listing of gene sets involved in Figure. 3C Venn diagram, comparisons of gene set cluster components in Nrn1$^{-/-}$ under resting vs. TCR restimulation condition.

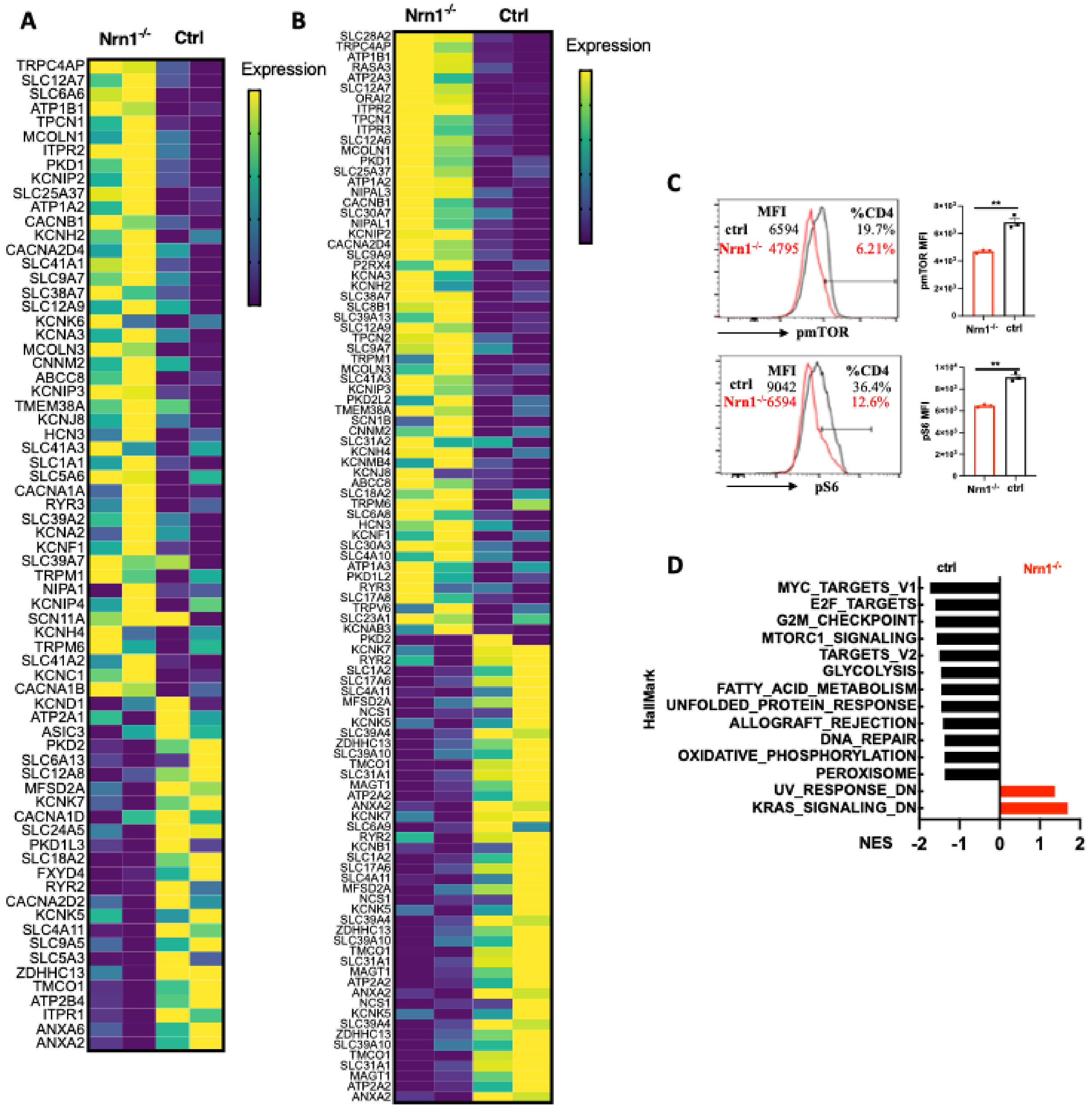


**Figure 3-figure supplement 1. Heatmap of differentially expressed genes and Hallmark gene set enrichment. (A)** Heatmap of differentially expressed genes in “GOMF_Metal ion transmembrane transporter activity” gene set from Nrn1$^{-/-}$ and ctrl iTreg cells cultured under the resting condition. **(B)** Heatmap of differentially expressed genes in “GOMF_Metal ion transmembrane transporter activity” gene set from reactivated Nrn1$^{-/-}$ and ctrl iTreg cells. **(C)** Detection of pmTOR and pS6 in Nrn1$^{-/-}$ and ctrl iTreg cells. Data represents three independent experiments. **p<0.01. Unpaired Student’s t-tests were performed. **(D).** Enrichment of Hallmark gene set in activated Nrn1$^{-/-}$ and ctrl iTreg cells (P<0.05, FDR q<0.25).

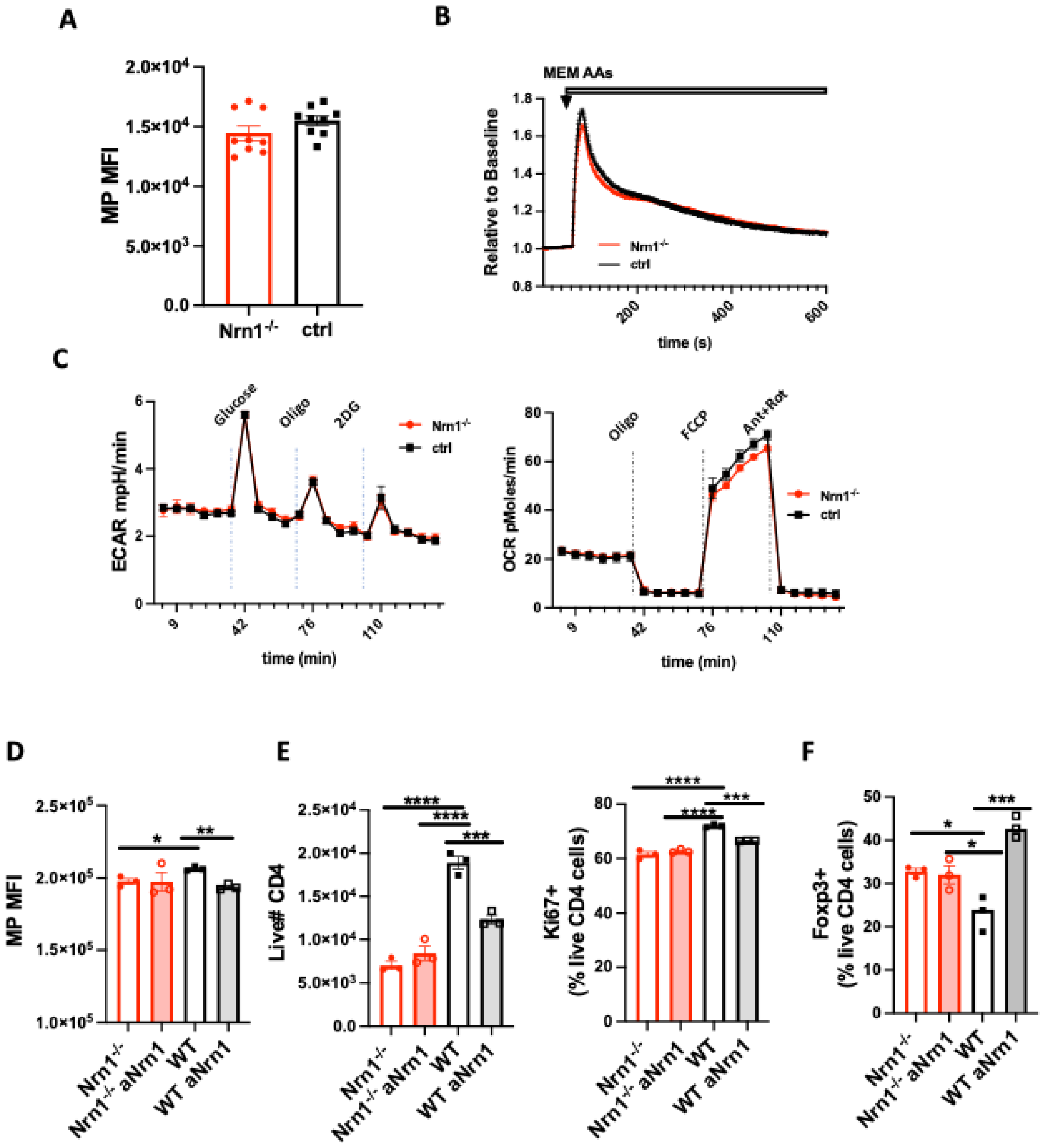


**Figure 3-figure supplement 2. Characterization of Nrn1$^{-/-}$ naïve CD4 T cells and effect of Nrn1 blockade on WT iTreg cell differentiation and expansion. (A)** Resting MP in Nrn1$^{-/-}$ and ctrl naïve CD4$^{+}$ T cells. **(B)** AAs induced MP change in Nrn1$^{-/-}$ and ctrl naïve CD4$^{+}$ T cells. **(C)** Seahorse analysis of extracellular acidification rate (ECAR) and oxygen consumption rate (OCR) in Nrn1$^{-/-}$ and ctrl iTreg cells. n=6~10 technical replicates per group. Data represent three independent experiments. **(D-F)** WT iTreg cells differentiated in the presence of Nrn1 antibody blockade. **(D).** MP in Nrn1$^{-/-}$ and WT iTreg cells differentiated in the presence or absence of anti-Nrn1 antibody. **(E).** Live cell number and proportion of Ki67 expressing cells after anti-CD3 restimulation among Nrn1$^{-/-}$ and WT iTreg cells differentiated in the presence or absence of anti-Nrn1 antibody. **(F).** Foxp3$^{+}$ cell proportion three days after anti-CD3 restimulation in Nrn1$^{-/-}$ and WT iTreg cells differentiated and restimulated in the presence or absence of anti-Nrn1 antibody. Data represent three independent experiments. *$p<0.05$, **$p<0.01$, ***$p<0.001$, ****$p<0.0001$. Ordinary one-way ANOVA was performed for multi-comparison.

**Figure 4-figure supplement Table 1: Gene sets enriched in Nrn1$^{-/-}$ and ctrl Te cells.**

| | |
|---|---|
| **Nrn1$^{-/-}$** | **Monoamine transport** |
| | IMPORT_INTO_CELL |
| | MONOAMINE_TRANSPORT |
| | CATECHOLAMINE_TRANSPORT |
| | NEUROTRANSMITTER_REUPTAKE |
| | **Membrane repolarization** |
| | CELL_CELL_SIGNALING_INVOLVED_IN_CARDIAC_CONDUCTION |
| | CARDIAC_MUSCLE_CELL_ACTION_POTENTIAL_INVOLVED_IN_CONTRACTION |
| | MEMBRANE_REPOLARIZATION |
| | CARDIAC_MUSCLE_CELL_MEMBRANE_REPOLARIZATION |
| | CARDIAC_MUSCLE_CELL_ACTION_POTENTIAL |
| | CARDIAC_CONDUCTION |
| | MEMBRANE_REPOLARIZATION_DURING_CARDIAC_MUSCLE_CELL_ACTION_POTENTIAL |
| | **Cell junction** |
| | CELL_CELL_JUNCTION_ASSEMBLY |
| | ADHERENS_JUNCTION_ORGANIZATION |
| **Ctrl** | **Transporter & Oxidoreduction** |
| | ACTIVE_TRANSMEMBRANE_TRANSPORTER_ACTIVITY |
| | PROTON_TRANSMEMBRANE_TRANSPORTER_ACTIVITY |
| | OXIDOREDUCTASE_ACTIVITY_ACTING_ON_NAD_P_H |
| | PRIMARY_ACTIVE_TRANSMEMBRANE_TRANSPORTER_ACTIVITY |
| | OXIDOREDUCTION_DRIVEN_ACTIVE_TRANSMEMBRANE_TRANSPORTER_ACTIVITY |
| | ELECTRON_TRANSFER_ACTIVITY |
| | **Sensory perception** |
| | SENSORY_PERCEPTION_OF_CHEMICAL_STIMULUS |
| | SENSORY_PERCEPTION_OF_SMELL |

Gene sets involved in Fig. 4F gene sets clusters enriched in Nrn1$^{-/-}$ and ctrl Te cells.

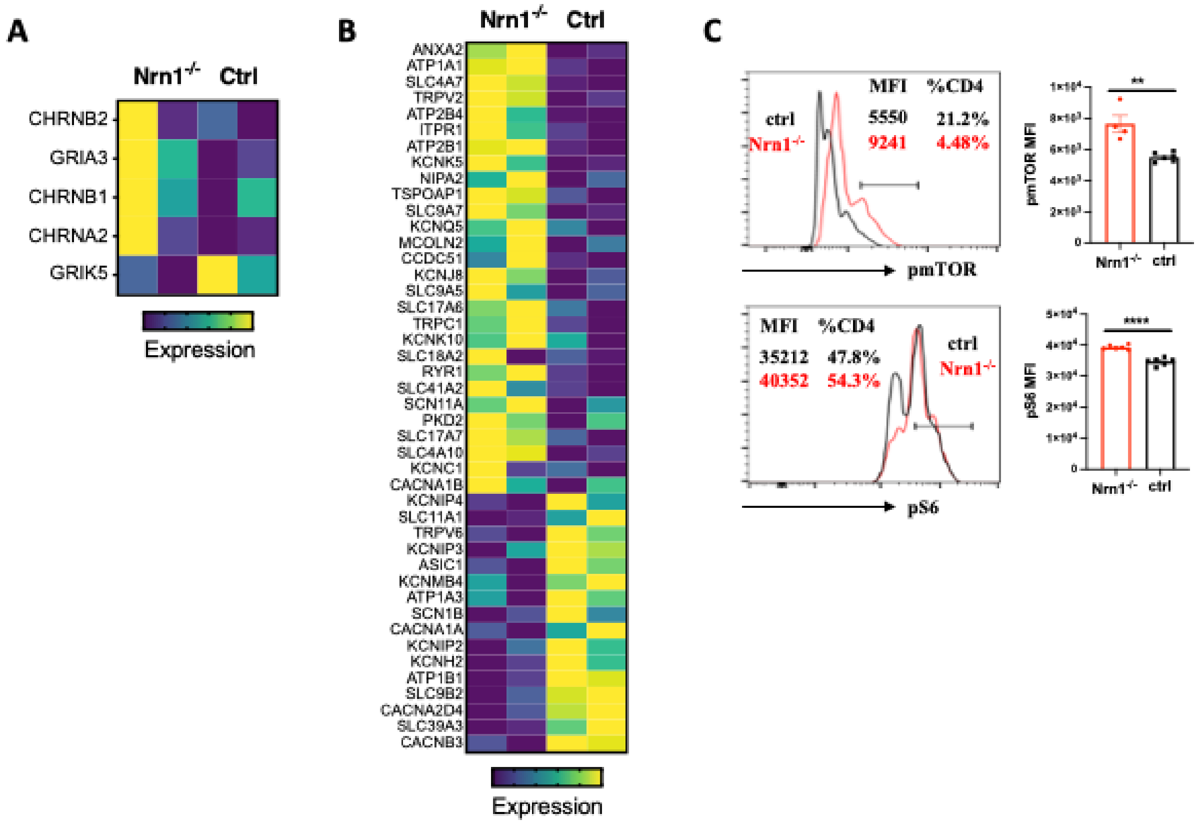


**Figure 4-figure supplement 1. Heatmap of enriched genes in Te cells. (A)** Differentially expressed genes in "GOMF_Neurotransmitter receptor activity involved in the regulation of postsynaptic membrane potential" gene set in Nrn1$^{-/-}$ and ctrl Te cells. **(B)** Heatmap of differentially expressed genes in "MF_metal ion transmembrane transporter activity" in Nrn1$^{-/-}$ and ctrl Te cells. **(C).** Detection of pmTOR and pS6 in Nrn1$^{-/-}$ and ctrl iTreg cells. Data represents three independent experiments. $^{**}p<0.01$, $^{****}p<0.0001$. Unpaired Student's t-tests were performed.

**Reference for supplement figure.**

Plitas, G., C. Konopacki, K. Wu, P.D. Bos, M. Morrow, E.V. Putintseva, D.M. Chudakov, and A.Y. Rudensky. 2016. Regulatory T Cells Exhibit Distinct Features in Human Breast Cancer. *Immunity* 45:1122-1134.
